# Single nucleus pituitary transcriptomic and epigenetic landscape reveals human stem cell heterogeneity with diverse regulatory mechanisms

**DOI:** 10.1101/2021.06.18.449034

**Authors:** Zidong Zhang, Michel Zamojski, Gregory R. Smith, Thea L. Willis, Val Yianni, Natalia Mendelev, Hanna Pincas, Nitish Seenarine, Mary Anne S. Amper, Mital Vasoya, Venugopalan D. Nair, Judith L. Turgeon, Daniel J. Bernard, Olga G. Troyanskaya, Cynthia L. Andoniadou, Stuart C. Sealfon, Frederique Ruf-Zamojski

## Abstract

Despite their importance in tissue homeostasis and renewal, human pituitary stem cells (PSCs) are incompletely characterized. We describe a human single nucleus (sn) RNAseq and ATACseq resource from pediatric, adult, and aged pituitaries (snpituitaryatlas.princeton.edu) and characterize cell type-specific gene expression and chromatin accessibility programs for all major pituitary cell lineages. We identify uncommitted PSCs, committing progenitor cells, and sex differences. Pseudotime trajectory analysis indicates that early life PSCs are distinct from the other age groups. Linear modeling of same-cell multiome data identifies regulatory domain accessibility sites and transcription factors (TFs) that are significantly associated with gene expression in PSCs compared to other cell types and within PSCs. Modeling the heterogeneous expression of two markers for committing cell lineages among PSCs shows significant correlation with regulatory domain accessibility for *GATA3*, but with TF expression for *POMC*. These findings characterize human stem cell lineages and reveal diverse mechanisms regulating key PSC genes.

## Introduction

Tissues are composed of several cell types that can assume different gene expression states in response to environmental cues^1^. Major objectives of current biological research include resolving cellular heterogeneity within tissues and elucidating the regulatory mechanisms determining cell types and states. With the recent development of single-cell (sc) omics technologies, researchers have refined the characterization of cell types in many tissues^2, 3^.

The pituitary gland secretes hormones that control crucial physiological processes, including reproduction, metabolism, and the stress response. The adenohypophysis represents the main portion of the pituitary gland and contains five hormone-producing cell lineages. Despite the physiological relevance of the pituitary in health and disease, human sc RNAseq studies to date have omitted the post-natal pituitary^4, 5^. Furthermore, mapping the pituitary epigenome landscape has not been included in the ENCODE project^6, 7^, and no chromatin accessibility profiling of the human pituitary at sc resolution has been reported.

Of particular interest is the insight into pituitary stem cells (PSCs) to be obtained from sc analyses. Pituitary hormone deficiencies, which include congenital hypopituitarism (combined pituitary hormone deficiencies), acquired hypopituitarism (secondary to trauma or surgery), as well as pituitary tumors such as adenomas, result in a severe disruption of endocrine systems and cause significant morbidity^8^. Thus, there is a need to develop stem cell therapies that could restore lost or damaged endocrine cell populations in the pituitary. Previous mouse studies demonstrated the existence of PSCs and their ability to self-renew and differentiate into all five endocrine cell types^9, 10^, thus opening potential therapeutic avenues for human pituitary deficiencies and pituitary tumors^11, 12^. Little is known about the epigenetic landscape and dynamics of human PSCs during post-natal life, which is critical information for realizing their therapeutic potential. Sc studies of human pituitary are important for resolving cell identities and revealing the regulatory mechanisms of this key cell type.

One impediment to characterizing human PSC heterogeneity and elucidating gene regulatory mechanisms through sc studies is the technical difficulty in generating high-quality datasets from the frozen *post-mortem* pituitary samples provided by tissue banks. We recently developed an integrated single nucleus (sn) multi-omics analysis using frozen adult murine pituitary^13^. Here, we successfully employed a similar procedure to characterize all major cell types in the human pituitary with a particular focus on PSCs. Archived frozen *post-mortem* pituitaries from pediatric, adult, and elderly subjects (one male and one female per age group) were jointly analyzed by snRNAseq and snATACseq (sn multi-omics). Importantly, we also generated a same-cell female pediatric pituitary sn multiome dataset. These analyses enabled us to characterize the transcriptome and chromatin accessibility landscapes of pituitary cell types. We further refined the identification of human PSC subtypes and their changes during aging and provide insight into the diverse gene regulatory mechanisms underlying stem cell identity and commitment.

## Results

### Sn multi-omics profiling of human pituitaries

To construct cell-type genome-wide maps of gene expression and open chromatin in the human pituitary, we conducted same-sample multi-omics assays of sn transcriptome (snRNAseq) and sn chromatin accessibility (snATACseq) in frozen *post-mortem* pituitaries from pediatric, adult, and aged individuals of both sexes that had been stored in tissue banks at −80 C for an average of ~10 years since donation (range 4 to 20; **Supplementary Table 1**). Because nuclei isolated from the same pituitary fragment were processed for both snRNAseq and snATACseq, the paired datasets, although not from the same nuclei, were sampled from the same population of nuclei in each pituitary studied. Additionally, to test for tissue heterogeneity and accuracy of cell type mapping across assays, and to improve inference of regulatory mechanisms, the remaining sample comprising nearly the entire pituitary from one female subject was pulverized and the isolated nuclei were then used to carry out: a) same-sample analysis of sn transcriptome and sn chromatin accessibility, b) same-cell sn multiome analysis providing simultaneous measurement of RNA expression and chromatin accessibility within each individual nucleus (**Fig. 1a**).

**Figure 1:**
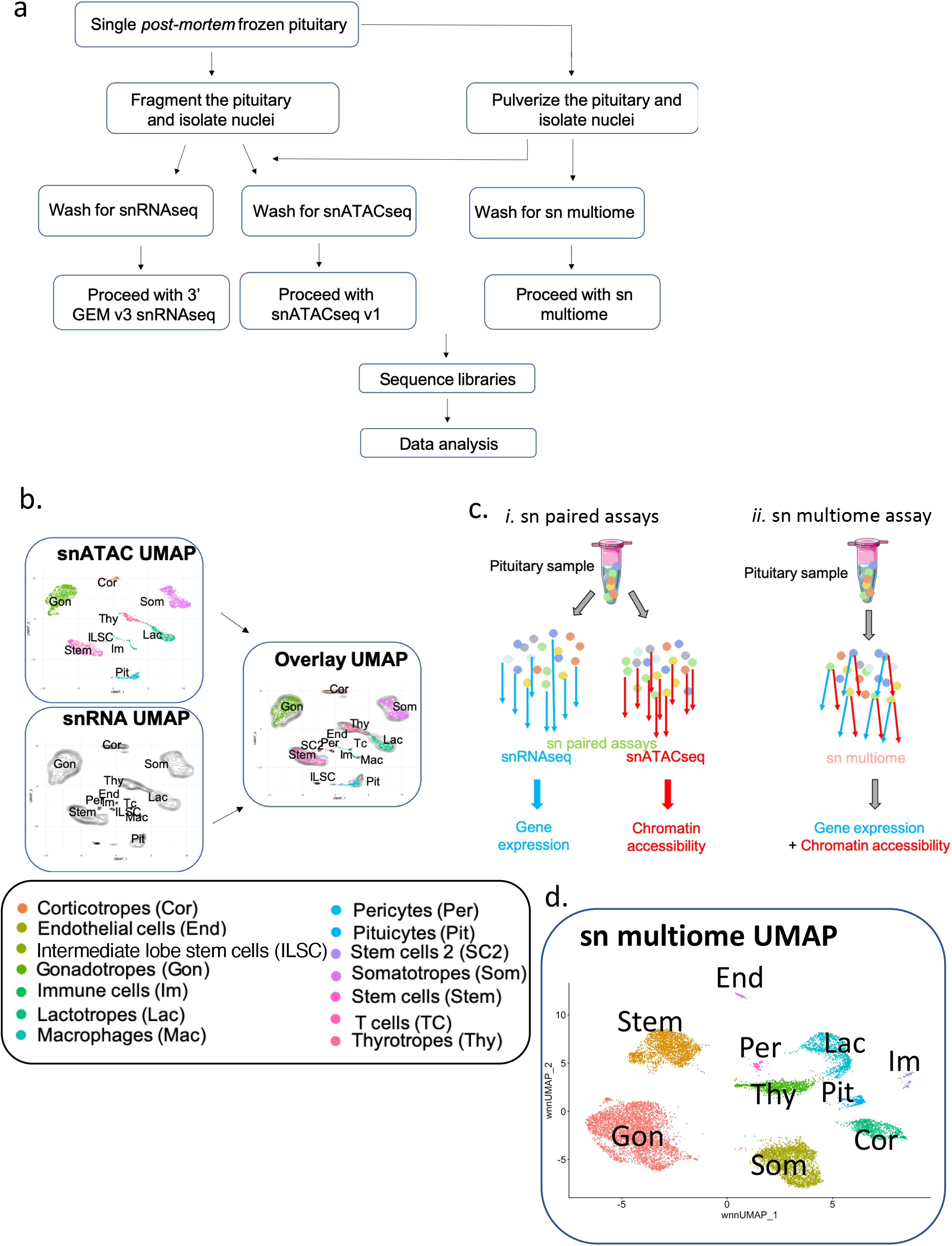
Experimental design for human pituitary cell type identification. **a**. Schematic of the overall experimental workflow, from procurement of the frozen pituitaries to sn data analysis. **b**. Schematic summarizing sn data integration. For each sample, the snATACseq dataset (colored dots UMAP) was integrated with the snRNAseq dataset (black contours UMAP) to generate an integrated multi-omics overlay UMAP identifying cell types. On the UMAP, cell types are color-coded and designated with a 2- to 3-letter code, as indicated on the bottom key. The female pediatric pituitary sample is represented as an example. All integrated samples are presented in **Supplementary Figs. 1** and **2**. **c.** Schematic of the comparison between sn paired assays (same-sample sn multi-omics) (*i*) and sn mutiome assay (same-cell) (*ii*). **d.** Same-cell sn multiome UMAP from the female pediatric sample (see **Supplementary Table 1**).

All snRNAseq and snATACseq libraries generated from the same samples were pooled for sequencing to reduce batch effects. Data meeting the quality control (QC) threshold were obtained from a total of 76,016 nuclei for snRNAseq and 44,141 nuclei for snATACseq in paired assays, and from 15,024 nuclei in the same-cell sn multiome assay (**Supplementary Tables 2 and 3**). For data analysis of a given sample processed through the same-sample sn paired assays, we generated UMAPs for both snATACseq and snRNAseq datasets, each identifying cell clusters by type (**Fig. 1b, Supplementary Fig. 1**). Integration of both datasets resulted in an overlay UMAP showing good correspondence of the major pituitary cell types across assay modalities (**Fig. 1b, Supplementary Fig. 2**). The same-cell sn multiome assay, in which each cell yielded both RNAseq and ATACseq datasets (**Fig. 1c**), directly generated an integrated UMAP plot (**Fig. 1d**).

### Transcriptome analysis of human pituitary cell types

The same-sample paired assay snRNAseq datasets had an average of 86% of reads mapped to the transcriptome and allowed for the detection of ~2,800 genes per nucleus, with comparable high-quality QC metrics obtained from all samples (**Supplementary Table 2a**). In the snRNAseq data analysis of individual male and female pituitary samples, cells were clustered using Seurat, visualized using t-SNE representation (**Supplementary Fig. 3**), as well as projected on UMAPs (**Supplementary Fig. 1**). Cell clusters were annotated manually using differential RNA expression of established pituitary marker genes. Key cell type markers included *FSHB*, *LHB*, and *GNRHR* for gonadotropes; *GH1* for somatotropes; *POMC* for corticotropes; *DIO2* for thyrotropes^5, 14^; *PRL* for lactotropes; and *SOX9*^10^, *LGR4*^15^, and *RBPMS* ^14^ for PSCs. A list of established markers used for the assignment of each cell type as well as new markers identified in our datasets are shown in **Supplementary Table 4**. The RNA counts (**Supplementary Fig. 4**), mitochondrial gene content (**Supplementary Fig. 5**), and ribosomal protein gene content (**Supplementary Fig. 6**) all indicated the high quality of the snRNAseq data obtained from each individual donor. Cell clustering analysis revealed well-defined cell clusters, including the five major hormone-producing cell types as well as several non-endocrine cell types (**Supplementary Fig.3**).

### Chromatin accessibility analysis of human pituitary cell types

The same-sample paired assay snATACseq datasets generated approximately 11,000 DNA fragments per nucleus with an average TSS enrichment score of 5.0 and a fraction of reads in called peak regions (FRiP) score of 47% (**Supplementary Table 2b**). Cells were clustered and visualized using UMAP representation (**Supplementary Figs. 1,2**). Cell clusters were manually annotated based on chromatin accessibility (i.e. peaks of accumulated reads) at informative promoters among the same marker genes used for the RNAseq annotation (see **Supplementary Table 4, Supplementary Fig. 7).** Thyrotrope cells were too poorly represented to generate reliable chromatin tracks, consistent with their being the lowest abundance endocrine cell type in the anterior pituitary^16^. Similar to the sn transcriptome analysis results, cell clustering of the snATACseq data from each donor resulted in distinct cell clusters with all the major cell types being identified, although a thyrotrope cluster could not be distinguished in all male samples (**Supplementary Figs. 1,2**).

### Cell type identification in snRNAseq and snATACseq datasets

Integration of the snRNAseq and snATACseq data from each sample was accomplished by label transfer from the snRNAseq to the snATACseq data using the Seurat pipeline (**Supplementary Figs. 1,2**). The major pituitary cell type clusters were detected in all individual samples. Some clusters showed a gradient of expression and chromatin accessibility, resulting in their distinction as separate clusters, although they were not physically distinct (lactotropes in **Supplementary Fig. 3d,** corticotropes in **Supplementary Fig. 3b** and **Supplementary Fig. 1f**; somatotropes in **Supplementary Fig. 3a,c,d** and **Supplementary Fig. 1d**; pituicytes in **Supplementary Fig. 3e,f;** gonadotropes in **Supplementary Fig. 3a,d**).

To improve the resolution of human pituitary cell types and to assess inter-individual variation, we merged same-sex snRNAseq datasets and color-labeled them by donor (**Fig. 2a,c**). Similarly, we merged the snATACseq data from same-sex samples, and labeled them by donor (**Fig. 2b,d**). In addition to the five endocrine pituitary cell types, we identified stem cells, pituicytes, as well as pericytes, endothelial cells, and immune cells (macrophages, T-cells, B-cells). We observed donor-to-donor heterogeneity in cell type clustering in both datasets. For example, in males, separate gonadotrope, somatotrope, and lactotrope clusters were noted in both RNAseq and ATACseq data, originating almost exclusively from the pediatric sample. In females, one gonadotrope, one somatotrope, and one stem cell cluster were also derived from the pediatric sample. The proportions of the major pituitary cell types identified by snRNAseq vs. snATACseq across all samples were highly correlated, indicating an agreement of major cell type assignment across the two assay modalities (R^2^ > 0.96; **Fig. 2e**). We also saw a similar distribution of the expression markers in human pituitary cell types in comparison with the same markers in an adult mouse pituitary dataset (^13^, **Supplementary Fig. 7**). A sn multiome dataset and a same-sample sn paired dataset were generated from the pulverized pediatric female sample, which further supported the reliability of cell type assignment across the two assays (**Supplementary Tables 3, 5**). All datasets are publicly available and are accessible for exploration at snpituitaryatlas.princeton.edu.

**Figure 2:**
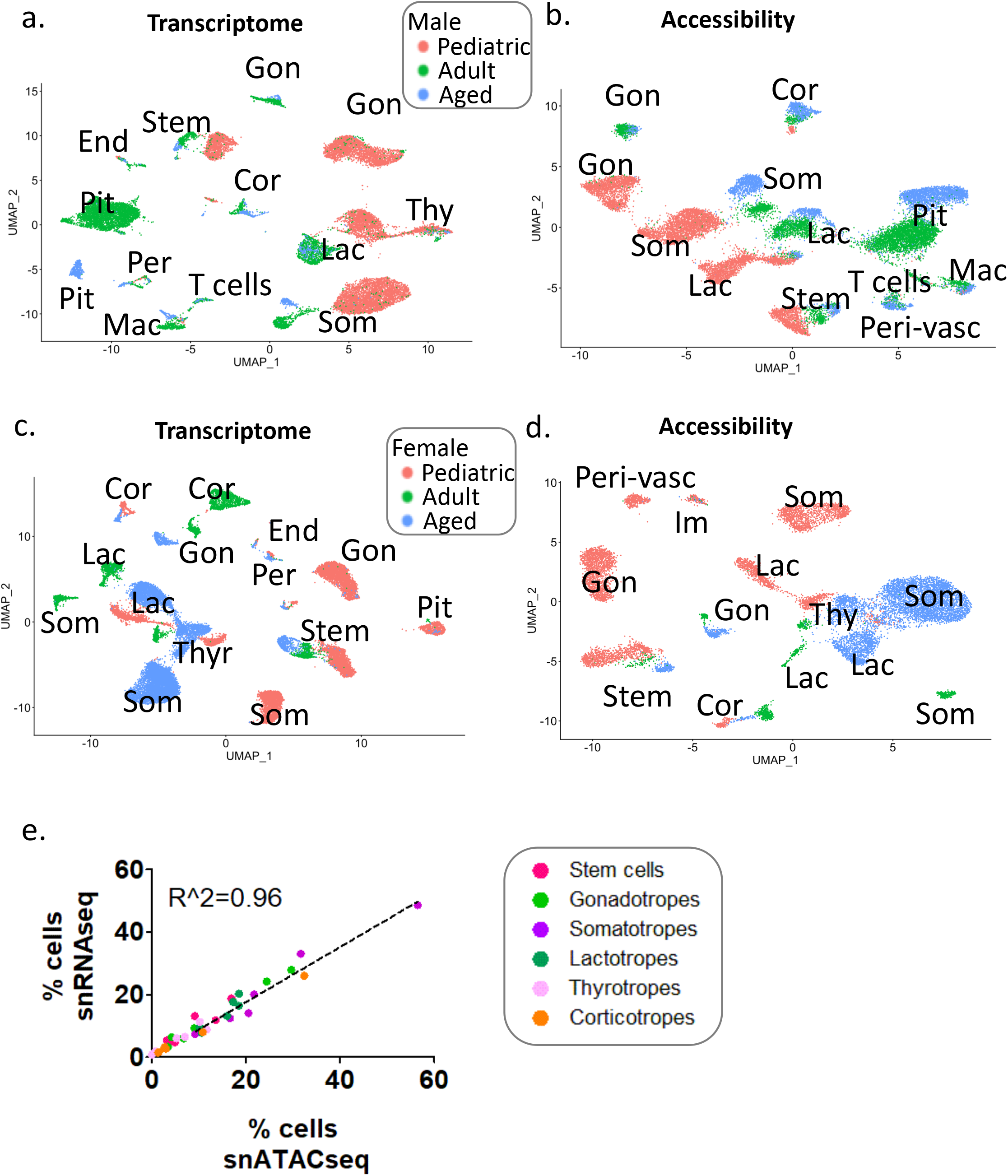
Merged analysis of same-sex human pituitary. **a-d**. t-SNE representation of sn transcript expression (**a**, males; **c**, females) and of sn chromatin accessibility (**b**, males; **d**, females) in the merged same-sex samples, with labeling by age of the subject in each sex. Individual subjects are color-coded as indicated. Each cluster is identified by a letter code as defined in **Fig. 1**. Donor-related information is provided in **Supplementary Table 1. e.** Correlation between the cell type proportions identified by snRNAseq vs. snATACseq for all samples (males and females). The linear regression is plotted. Pituitary cell types are color-coded and the key is provided on the right.

### Characterization of the PSC population

The stem cells from all samples identified by snRNAseq (**Fig. 2**) were re-clustered using the Seurat pipeline, leading to the detection of 9 clusters, 6 of which were not well separated and formed a large group (**Fig. 3a**). Three clusters that are highlighted in **Fig. 3a** corresponded to lineage-committed progenitor stem cells, with each distinguished by *POMC*, *POU1F1*, and *GATA3* expression, respectively (see Discussion). The remaining large group of 6 clusters expressed *SOX2*, *SOX9*, and the Hippo pathway effectors *WWTR1* (a.k.a. *TAZ*) and *YAP1* (^17^ and reviewed in ^18^; **Fig. 3d** and **Supplementary Fig. 8**), which are indicative of uncommitted PSCs, with cluster 5 showing the highest expression for some of these markers.

**Figure 3:**
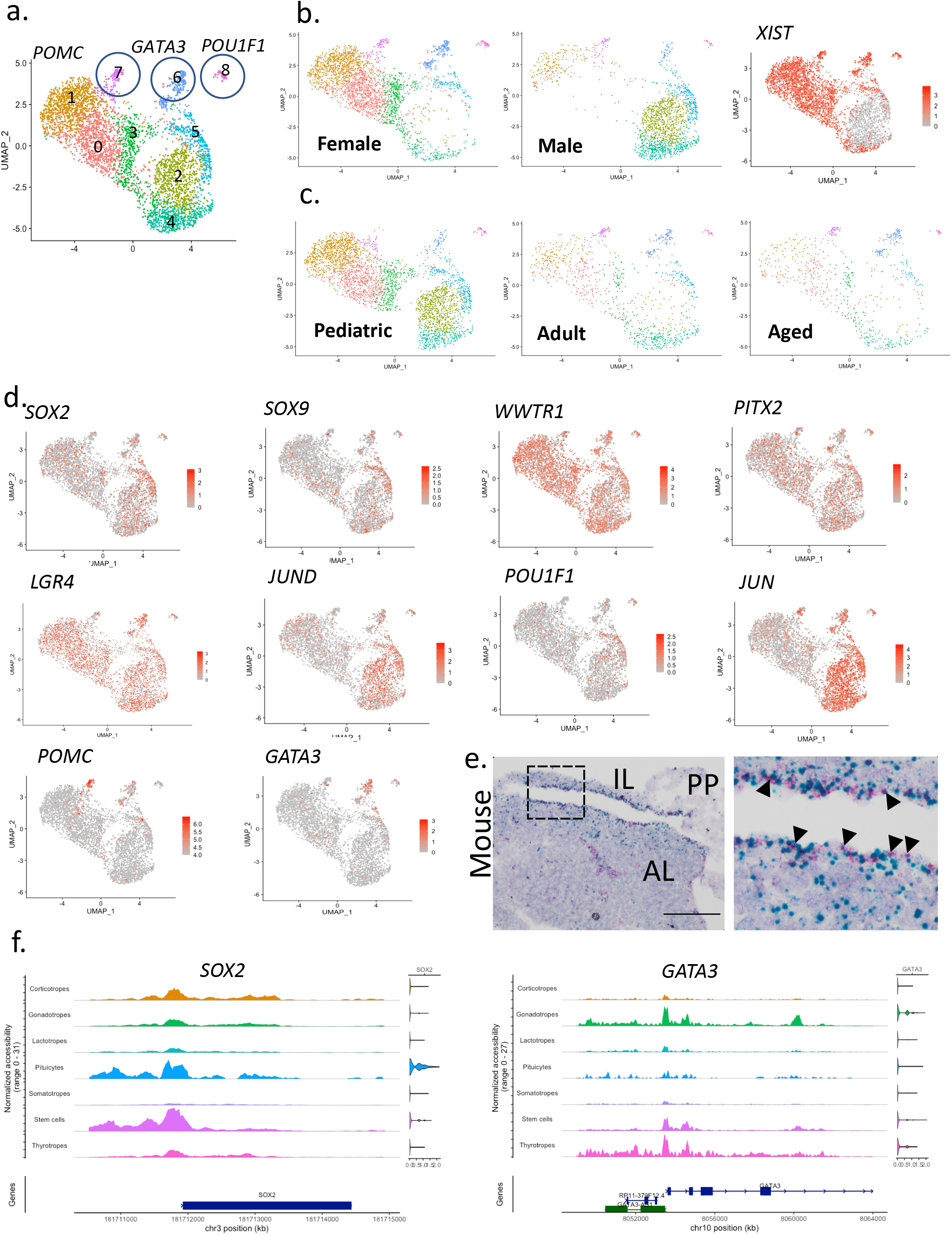
Identification of human stem cell sub-clusters by snRNAseq. **a**. UMAP showing the stem cell cluster identification based on the snRNAseq data from the six merged human pituitary samples. Each cell cluster is color-coded and numbered. Lineage-committed progenitor stem cells are circled. **b**. UMAPs identifying all color-coded stem cell sub-clusters in females and males. The feature plot on the right shows *XIST* expression, highlighting the female samples. **c**. UMAPs identifying all color-coded stem cell sub-clusters in the pediatric, adult, and aged subjects. **d.** Feature plots depicting the expression distribution of key stem cell marker genes and of cell lineage commitment marker genes among the various clusters. A scale is included for each feature plot. All scales are similar except for *POMC* due to background gene expression. Additional gene feature plots are presented in **Supplementary Fig. 8**. **e**. Colocalization of *Sox2* (red) and *Jun* (blue) transcripts in a wild-type P56 CD-1 male adult mouse pituitary. Scale bar is 200μm. AL: anterior lobe; IL: intermediate lobe; PP: posterior pituitary. *Left*, full image. *Right*, magnification of the boxed region in the left panel. Arrows highlight specific cells with colocalization of *Sox2* and *Jun*. Refer to **Supplementary Fig. 10** for *Sox2* and *Jun* colocalization at P3 and P15. f. Gene expression analysis (violin plots at the right of each figure) and chromatin accessibility tracks analysis for *SOX2 (Left*) and *GATA3 (Right*) in all pituitary cell types from the sn multiome dataset generated from the pediatric female. The gene structure is presented below the tracks.

We next examined the relationship of the PSC clusters to the sex and age of the donors (**Fig. 3b,c**). The uncommitted stem cell clusters were largely separated in samples from each sex, as confirmed by expression of the female-specific *XIST*^19^. When the male and female datasets were grouped by age, all PSC subtypes were represented at all ages studied. The distribution of RNA markers for progenitor and committing stem cells is shown in **Fig. 3d**. The canonical stem cell markers *SOX2* and *SOX9*, as well as genes previously implicated in pituitary stem cell regulation (i.e. *WWTR1*, *PITX2*, and *LGR4*; for review, see ^18^), were broadly expressed. Interestingly, expression of *JUN* and *JUND*, which were implicated in the regulation of stemness in other tissues^20, 21^, was heterogeneous, with the highest expression associated with clusters that were predominant in male samples.

We also compared the patterns of marker gene expression in human and mouse PSCs. As expected, we detected *Sox2*, *Sox9*, *Wwtr1*, and *Yap1* across the stem cell population of both species (**Fig. 3d; Supplementary Figs. 8,9**). Expression of *POU1F1* in human samples was detected in a proportion of the cells in the main PSC cluster, indicating committing progenitors amongst this population, as seen for *Pou1f1* in mouse, and similarly for *PAX7/Pax7* (a determinant of intermediate lobe and melanotrope identity; ^22^). Additional markers that were either previously reported in mouse pituitary stem cells or linked to stemness, were also found in human stem cells, including *WIFI*^23, 24^, *HES1*, *NOTCH2*^18^, *SMAD4*^25^, and *SMAD5*^26, 27^ (**Supplementary Fig. 8**).

*JUN*, which was expressed in uncommitted, predominantly human male PSCs, has not been proposed as a PSC marker but was previously reported to be enriched in SOX2 positive cells through bulk sequencing in mouse^15^. We therefore examined whether *Jun* showed co-expression with the stem cell marker *Sox2* by *in situ* hybridization in neonatal, juvenile male, and adult male mouse pituitaries. This analysis confirmed *Jun* as a stem cell marker by identifying *Jun-Sox2* double labeling in samples from all ages (**Fig. 3e** and **Supplementary Fig. 10**). Overall, characterization of the heterogeneity of the PSC population from human and mouse supports the existence of different subtypes of uncommitted stem cells that are distinguishable from early committing lineages (see ^28^).

The acquisition of both snRNAseq and snATACseq data from the same samples provides high resolution analysis of the chromatin accessibility pattern of key genes within each pituitary cell type and reveals potential novel regulatory domains (see ^13^). Shown in **Fig. 3f** are cell type-specific gene expression and chromatin accessibility by cell type for the stemness marker *SOX2* and the putative gonadotrope/thyrotrope committing cell lineage marker *GATA3. SOX2* was expressed in stem cells and pituicytes, which both showed the highest promoter accessibility. *GATA3* was expressed in gonadotropes and thyrotropes in addition to the committing stem cell lineage. All three cell types also all showed more chromatin accessibility in the region of the *GATA3* promoter. Other *SOX2* and *GATA3* putative cis-regulatory domains showing increased accessibility in cell types showing expression are also evident. In a subsequent section, we elucidate further the regulatory control of these and additional key PSC markers by modeling same-cell sn multiome data.

### Diversity of PSC epigenetic programs

We next studied coordinated gene expression and chromatin accessibility programs in PSCs. We utilized the Pathway Level Information ExtractoR framework (PLIER) that deconvolves datasets into co-varying latent variable (LV) gene sets using known pathways, while not enforcing the strict orthogonality required for principal component analysis^29^. These PLIER analyses identified both RNA and chromatin accessibility LVs that were preferentially expressed in each major pituitary cell type (**Supplementary Figs. 11,12**). One RNA LV was stem cell-specific and highly expressed in both sexes at all ages (**Fig. 4a** for the top 30 genes, and **Fig. 4b** for the top 200 genes). Projection of this PSC LV onto adult mouse pituitary snRNAseq data^13^ showed conservation of this PSC transcriptome program in mouse (**Fig. 4c**). To determine whether this program was also associated with altered chromatin structure in PSCs, we projected this LV onto the human snATACseq data using the promoter accessibility signals as the gene features. This projection showed that the LV transcriptome program was associated with increased chromatin accessibility at the corresponding gene promoters (**Fig. 4d**). Analysis of the snATACseq data identified a number of largely distinct accessibility programs that were each most strongly activated in subjects of different ages or sex (**Supplementary Fig. 12**). The complexity of PSC chromatin programs identified in this analysis may be related to the diversity of the donors (see Discussion).

**Figure 4:**
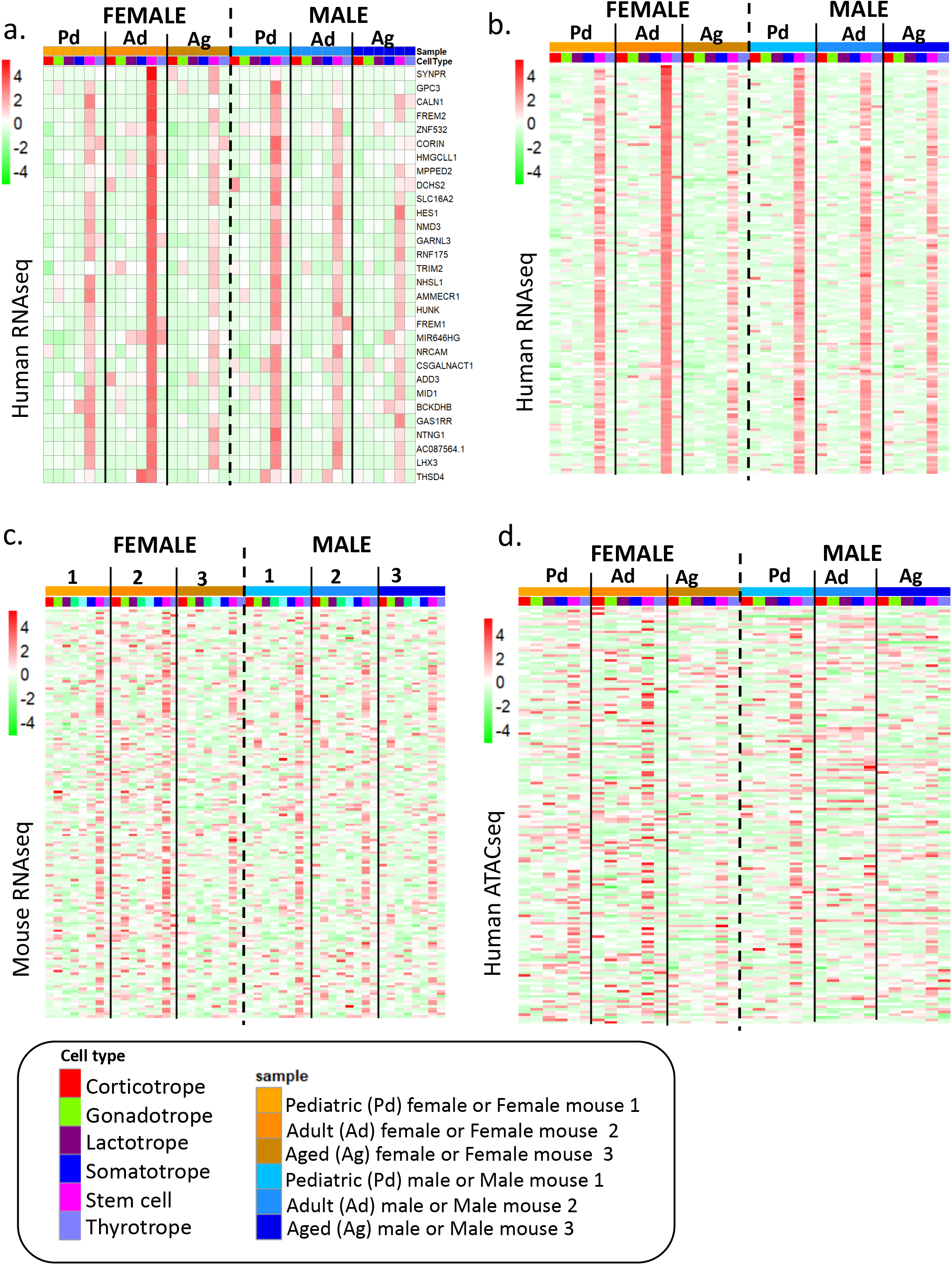
Characterization of coordinated gene expression and chromatin accessibility programs in human pituitary cell types. **a, b**. Heatmap of the levels of gene expression for the top human PSC LV (LVsc_rna_) for each cell type and donor, top 30 genes are shown in (**a**) and top 200 genes in (**b**). Each pituitary sample is indicated at the top. In the scale bars, red signifies the highest level of RNA expression or chromatin accessibility. Pd, pediatric; Ad, adult; Ag, aged pituitary. **c.** Heatmap showing the top 200 genes associated with the human LVsc_rna_ applied to the murine snRNAseq dataset^13^. **d.** Heatmap showing the top 200 genes associated with the human LVsc_rna_ applied to the human snATACseq datasets. (**a**, **b**, **c**, **d)** Cell type and subject color-coding are provided on the bottom key. Refer to **Supplementary Table 1** for donor-related information. Additional LV analyses are presented in **Supplementary Figs. 11 and 12**.

In addition to cell type-specific LVs, we also identified one chromatin accessibility LV showing significantly greater activity with increasing donor age (LVage_atac_). Shown in **Fig. 4e** is a heat map of the 30 highest weighted promoters comprising this LV in each sample by cell type. When the level of activity in each cell type was plotted separately across the age range studied by sex, we found that all cell types showed an increase in accessibility of these promoters with age, especially between the pediatric and adult samples (**Fig. 5a**). Notably, the increases in accessibility in the PSCs were less pronounced in females, while almost no changes were observed in males (pink lines in **Fig. 5a**.) These results suggest that this accessibility program represents age-associated coordinated changes that are more prominent in differentiated cells than in PSCs.

**Figure 5:**
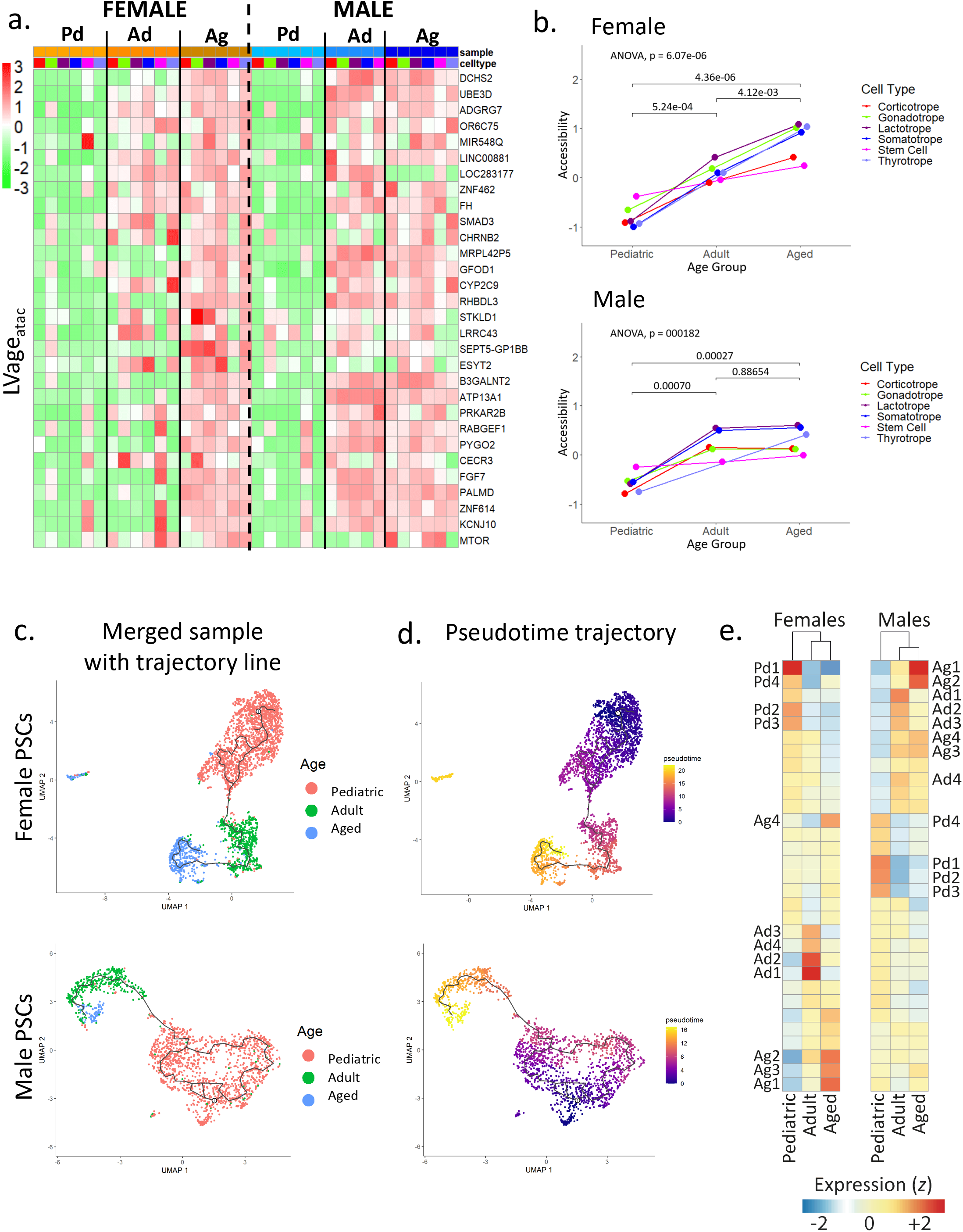
Age-associated chromatin accessibility and transcriptome pseudotime trajectory analysis. **a.** Heatmap showing the chromatin accessibility levels of the top 30 genes in the human age-dependent LV (LVage_atac_). Cell type and subject color-coding are provided on the bottom of **Fig. 4**. Refer to **Supplementary Table 1** for donor-related information. Additional LV analyses are presented in **Supplementary Figs. 11** and **12**. **b.** Plot showing the overall changes in chromatin accessibility for all pituitary cell types over age for the females (*Left*) and the males (*Right*). Pituitary cell types are color-coded. The same cell types are linked with lines over age of the subjects. **c**. UMAP showing the trajectory within the stem cell cluster with samples color-coded by age for female (*Top*) and male (*Bottom*) samples. **d.** Pseudotime trajectory analysis for the female (*Top*) and male (*Bottom*) samples. The trajectories from each starting point head to the older samples. The color scale is shown for the pseudotime trajectories. **e.** Gene modules identified with pseudotime and showing changes over age. Monocle 3 identified groups of genes that change over as a function of pseudotime per sex. Trajectory variable genes were grouped into modules which were then plotted on a heatmap to show the relative expression of each gene module, within a sex, in each age group. The top 4 enriched modules for each age group are labelled, with module 1 being the most highly enriched in each age. Pd1-4 indicates the Pediatric top 4 modules, Ad1-4 represents the Adult enriched modules, and Ag1-4 marks the Aged samples top 4 modules. Blue-red on the color scale represents low-high relative expression levels (z-transformed mean expression) of gene modules. See **Supplementary Fig. 13** for selected gene trajectories within specific modules. See **Supplementary Table 6** for the top genes per modules.

To further explore the relationship of PSC transcriptomes in samples from different ages, we constructed a pseudotime trajectory from same-sex snRNAseq datasets using the Monocle algorithm^30^ (**Fig. 5b,c**). In females as well as in males, the region of the graph most densely occupied by the pediatric PSCs was chosen as the root of the trajectory. In both sexes, pediatric PSCs formed the largest group, which separated from the adult and aged PSCs. This separation shows the large differences between PSC transcriptomes from pediatric and adult samples and also suggests that we have not captured all transitional stages of stem cells in the samples analyzed. To specify sets of genes that are dynamically regulated as cells progress along the trajectory, we identified several correlated gene modules per age group in females and in males (**Fig. 5d**). The top genes in the most significant modules and the trajectories of selected genes are shown in **Supplementary Table 6** and **Supplementary Fig. 13**, respectively. Notably, almost none of the module-defining transcripts were previously reported as PSC markers, and their roles in PSC physiology over the lifespan are not known. Overall, these analyses show the relative stability, an aging-related chromatin program in PSCs, and dynamic changes in the PSC transcriptome with aging.

### Transcription factor and epigenetic control mechanisms of PSC genes

Using snRNAseq and snATACseq datasets obtained from the same mouse pituitaries, we recently reported that chromatin accessibility is a key determinant for cell type transcriptional programs^13^. In comparison to same-sample datasets, same-cell sn multiome data confer vastly greater statistical power for inferring the regulatory mechanisms underlying expression of specific genes^31, 32^. The matched transcriptome and chromatin accessibility data in same-cell sn multiome assays allow the co-variation of chromatin accessibility and gene expression to be modeled in thousands of individual cells. Additionally, not all cells within a cell type express the same transcripts. Same-cell sn multiome data have the potential, for the first time, to provide insight into transcription factors (TFs) and epigenetic mechanisms that shape heterogenous gene expression within the same pituitary cell type.

To explore the role of alterations in TF expression and chromatin state in modulating key PSC genes, we applied a linear modeling computational framework to the 15,024 nuclei in the same-cell sn multiome dataset obtained from a pediatric female pituitary. For each target gene, the linear model selects potential cis-regulatory regions comprising ATAC promoter peaks as well as co-accessible distal peaks. Then, testing the co-expression of putative trans-acting regulatory factors that have predicted TF binding sites in the co-accessible regions, linear regression identifies the TFs and chromatin regions most significantly predictive of expression of the target gene. The linear model, when used to analyze all pituitary cells (“pan pituitary cell” analysis), can infer the mechanisms and factors implicated in cell type-specific expression. When only cells comprising one pituitary cell type are analyzed, the linear model can generate hypotheses for the mechanisms responsible for differential expression among the different cells comprising this lineage.

We first analyzed the committed progenitor markers *POMC*, *POU1F1*, *TBX19*, and *NR5A1*. The output of the linear model pipeline is the p-value that each selected cis-regulatory region and each individual TF with sites in that region contribute to the expression of the target gene (**Supplementary Figs. 14-17**). The TFs contributing to cell type-specific expression of these marker genes using pan pituitary cell analysis included many factors that were previously implicated in the differentiation of committing stem cells. For example, the *POMC* analysis identified TBX19, which is an inducer of the *POMC-expressing* corticotrope/melanotrope lineage and of *POMC* expression^33^. *TCF7L2*, which was highly significant in the analyses of *POU1F1* and *TBX19*, is an effector of the WNT signaling pathway, which regulates pituitary growth and development^34^. Similarly, *LEF1*, another mediator of WNT signaling (see ^35^), was also identified in the *TBX19* analysis. ESR1 (estrogen receptor alpha) was the most significant TF implicated in *NR5A1* expression. Consonant with this finding, a recent study in murine gonadotrope cell lines demonstrated that estrogen-dependent binding of this nuclear receptor to a newly identified enhancer region triggers *Nr5a1* expression during gonadotrope lineage specification^36^. The high significance obtained in the linear model analysis for transcriptional regulators that were reported in previous research suggests that new candidates we identified warrant consideration for future study. For example, the TF showing the second highest significance in the pan pituitary cell analysis of *POMC* is MNX1, an important homeobox gene previously implicated in motor neuron, pancreas, and lymphoid cell development^37, 38, 39^. Therefore, MNX1 is an intriguing new candidate transcriptional regulator in the commitment towards the corticotrope/melanotrope lineages. In addition to the identification of novel putative TF regulators, when applied to all pituitary cells, the model also specifies the proximal and distal regulatory sites significantly associated with expression of the target gene in the cell types expressing that gene. These analyses identify previously unexplored regulatory domains in these key PSC genes that show accessibility associated with gene expression and are therefore cis-regulatory domain candidates (**Fig. 6** and **Supplementary Figs. 14-18**).

**Figure 6:**
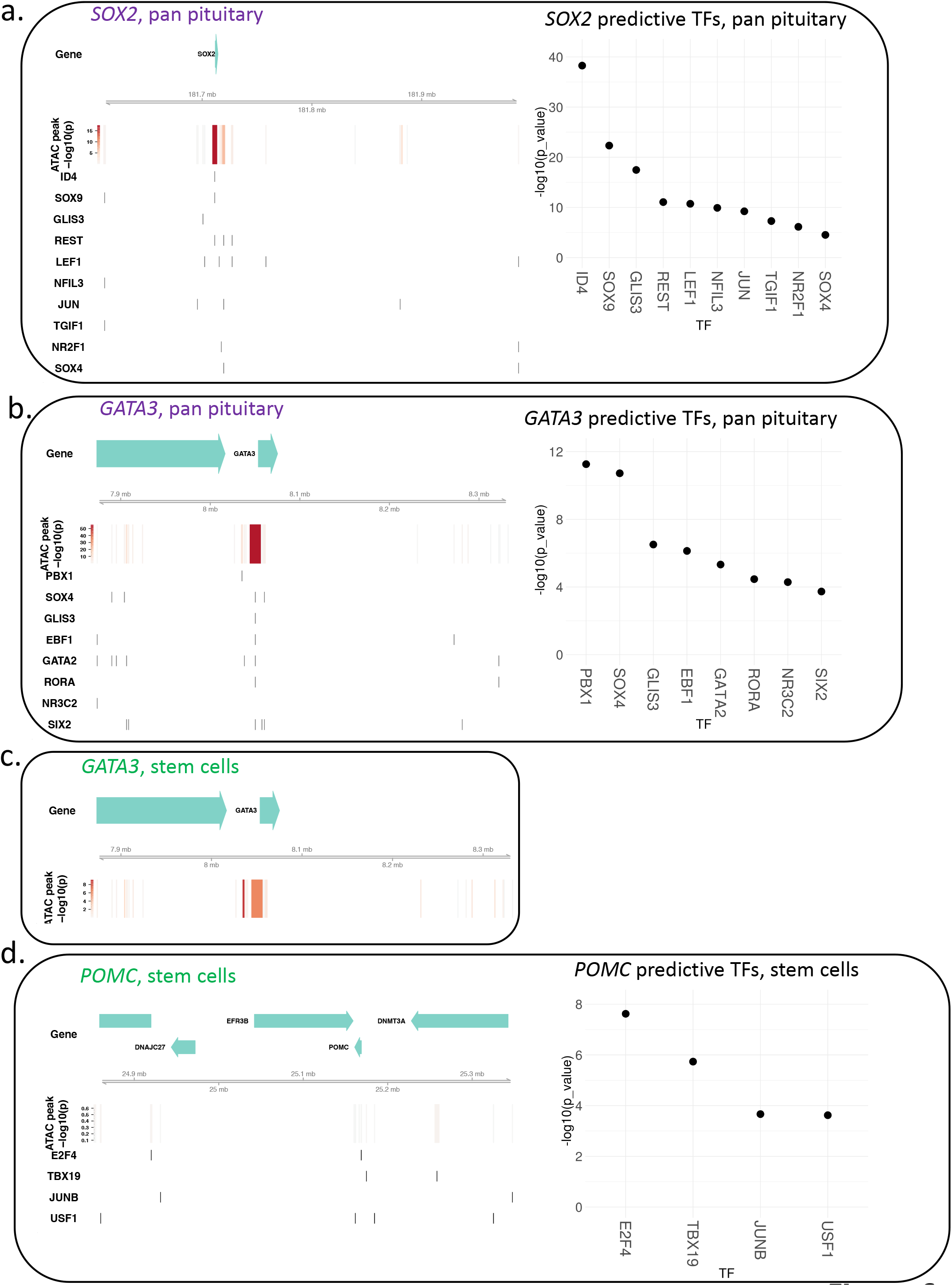
Linear model predicting the chromatin accessibility mechanisms and TFs contributing to PSC gene expression. **a**. Linear modeling analysis of all pituitary cells (“pan pituitary cell”) infers the chromatin accessibility and the TFs involved in stem cell-specific *SOX2* expression (*Left*, top track is the contribution of each peak to gene expression measured by −log(P-value), and bottom tracks are the TFs binding sites). The individual contribution of each predicted TF to *SOX2* expression is shown as −log(P-values, *Right*) See **Supplementary Fig. 18** for *SOX2* analysis in stem cells only. **b**. Pan pituitary cell analysis infers the chromatin accessibility and the TFs involved in stem cell-specific *GATA3* expression (*Top*, top track is the contribution of each peak to gene expression measured by −log(P-value), and bottom tracks are the TFs binding sites). The individual contribution of each predicted TF to *GATA3* expression is shown as −log(P-values) (*Right*) **c**. Linear modeling analysis in stem cells only infers the chromatin accessibility and the TFs involved in the differential expression of *GATA3* expression within the stem cell population. No TFs are predicted to contribute to *GATA3* expression in stem cells. **d**. Linear modeling analysis in stem cells only infers the chromatin accessibility and the TFs involved in the differential expression of *POMC* expression (*Left*, top track is the contribution of each peak to gene expression measured by −log(P-value), and bottom tracks are the TFs binding sites). The individual contribution of each predicted TF to *POMC* expression is shown as log(P-values) (*Right*). See **Supplementary Fig. 17** for pan pituitary cell analysis of *POMC* expression.

We next studied the stemness marker *SOX2* and the committing cell lineage marker *GATA3*. When all pituitary cells were examined, the expression of *SOX2* within the overall stem cell subtype was associated with highly significant co-accessible proximal, upstream, and downstream regulatory domains (**Fig. 6a,** *Left*) as well as expression of TFs mapping to these domains (**Fig. 6a,** *Right*). These results indicate that PSC-specific expression of *SOX2* depends on a pattern of chromatin accessibility of regulatory domains present within these cells as well as expression of the requisite regulatory factors interacting with these domains. A contrasting result was obtained when applying the linear model to only PSCs to infer the regulatory circuits involved in heterogeneous expression of *SOX2* within PSCs. In this analysis, cis-regulatory domains correlated poorly with *SOX2* expression (**Supplementary Fig. 18,** *Left*), and a restricted set of regulatory factors (**Supplementary Fig. 18**, *Right*) was implicated in the heterogeneous pattern of *SOX2* expression in PSCs. These results suggest that the chromatin structure is sufficient for *SOX2* expression in all PSCs and the expression within specific PSCs depends on the expression of key regulatory TFs. When performing a pan pituitary cell analysis for *GATA3*, we observed a pattern consonant with that of *SOX2*, with both chromatin structure and regulatory factor expression being responsible for expression in PSCs (**Fig. 6b**, *Left*, **Fig. 6b**, *Right*). However, contrary to *SOX2*, heterogenous *GATA3* expression within PSCs was associated with cis-regulatory chromatin accessibility domains, but not with expression of specific regulatory factors (**Fig. 6c**). These results suggest that the regulatory proteins needed for *GATA3* expression in PSCs were expressed in all of the cells, and the heterogeneous expression pattern within PSCs was determined by differences in chromatin accessibility of regulatory domains between *GATA3*-expressing and non-expressing PSCs. When *POMC* was analyzed only in PSCs, the most significant TFs identified were E2F4^40^ and TBX19, while the co-accessible regulatory regions were of low significance (**Fig. 6d**). These results suggest that differential *POMC* expression in committing PSCs vs. uncommitted PSCs is due to expression of these key TFs more so than to alterations in chromatin accessibility at key regulatory regions.

## Discussion

We generated high quality snRNAseq and snATACseq datasets from individual human pituitaries, demonstrating the feasibility of sn profiling in frozen *post-mortem* samples that had been stored at −80 C for as long as two decades. Analysis of these data provides insight into the heterogeneity of PSCs and the regulatory mechanisms and circuits underlying the expression of key PSC genes. We distinguish and characterize uncommitting and committing stem cell lineages, differences related to the age and the sex of the donors, and propose diverse mechanisms responsible for expression of key PSC markers.

Reclustering of the stem cells identified by snRNAseq data analysis distinguishes three clusters consistent with committing stem cells (see **Fig. 3a**). The *POMC-expressing* cluster is likely a precursor of the corticotrope/melanotrope lineages^41^. Another cluster expressing *POU1F1* represents PSCs with the potential to commit to the somatotrope, lactotrope, and thyrotrope lineages^42^. The *GATA3*-expressing cluster presumably comprises cells that are committing to the gonadotrope lineage, although low expression of *NR5A1* precludes definitive cell type lineage assignment as *GATA3* is also reported in thyrotropes (**Supplementary Fig. 8**)^43^. All three committing lineages were identified in both sexes. The uncommitted stem cells, which formed six clusters that were not well separated, were for the most part non-overlapping in male vs. female samples.

Our analysis identifies PSC transcriptome and epigenetic programs as well as age-related differences in PSCs. We identify one RNAseq PSC LV program that is well expressed in all samples assayed, conserved in mouse and associated with PSC-specific chromatin changes at the promoters for the genes comprising this LV. We also identify an ATACseq LV that exhibits increased accessibility with age in all pituitary cell types but shows smaller changes with age in PSCs. When the snRNAseq data were analyzed by trajectory, PSCs from the pediatric samples were separated from the adult and aged samples in both sexes. We find that human and mouse PSCs shared similar patterns of gene expression and were characterized by the presence of several subtypes of uncommitted and committed cells, suggesting a high degree of conservation of this cell type in evolutionary time. However, the identification of species-specific genes might signify inter-species or age-related differences, with potential implications for using the mouse model for therapy development.

The PSC LV programs, and the complete separation of PSCs across different ages in the trajectory analysis, indicate that fully characterizing the changes in PSC transcriptional and epigenetic programs with aging will require analysis of additional samples over the age span. Collectively, our data suggest that sex and age influence several biological processes in stem cells. Additional study is warranted to further elucidate the sex and age differences that are likely to have a significant impact on the development of new stem cell-based therapies.

The datasets generated encompass all major cell types in the human pituitary. Reliability of the identification of all major cell types in the same-sample snRNAseq and snATACseq datasets from both sexes and from a range of subject ages is supported by the concordance of cell type proportions obtained by both assays and the confirmation of cell type identification in the same-cell sn multiome data. We report gene expression and chromatin accessibility LVs that are characteristic of each major pituitary cell type. Extensive data from the female pediatric pituitary are provided by multiple same-sample datasets and a large same-cell sn multiome dataset. These data represent a resource to address questions about the characterization and regulatory mechanisms of any cell type in the human pituitary.

Inferences from the same-cell sn multiome dataset using the linear model provides insight into the TFs and accessible chromatin sites contributing to the expression of key PSC genes. The model also provides insight into the general mechanisms (TF expression, chromatin accessibility differences or both) responsible for differential expression of the target PSC genes among different cells. Because the model is based on detection of regulatory feature correlation with target gene expression, the results obtained when PSC genes are analyzed among all pituitary cells represent mechanisms implicated in target gene expression in PSCs in comparison with other cell types. When the model is applied only to PSCs, the results represent hypotheses for differential expression of these target genes within a subset of PSCs. For the PSC and committed progenitor markers analyzed (*POMC*, *POU1F1*, *TBX19*, *NR5A1*, *SOX2*, and *GATA3*), multiple chromatin accessibility sites and TFs predicted to bind to accessible sites are identified with high probability as contributing to stem cell expression of these markers in comparison with other cell types. This supports the formulation that the expression of each of these markers in PSCs depends on epigenetic remodeling of chromatin as well as on expression of key TFs that are necessary for driving gene expression.

Analysis of same-cell sn multiome data from the pediatric female pituitary suggests that a diversity of mechanisms contribute to differential expression of marker genes among PSCs. With respect to differential expression of PSC markers within PSCs, *NR5A1* and *POU1F1* show TFs and chromatin sites associated with heterogeneous expression. *POMC*, *TBX19*, and *SOX2* are associated with the expression of specific TFs and *GATA3* with accessibility of specific regulatory sites. The pan-pituitary analysis shows the importance of both TF expression and chromatin structure in the expression of key PSC genes. However, these analyses of heterogenous expression within PSCs suggest that the differential expression of some markers is predominantly determined by expression of key TFs in those cells, whereas the differential expression of other markers depends on heterogeneity in chromatin structure.

A strength of this study is the multi-omic profiling of the entire pituitary at postnatal stages, that we use to develop a map of the variation of PSCs between the sexes and with development and aging. Additionally, to our knowledge, this is the first study to profile human PSCs through ageing and to propose mechanisms for differential marker gene expression within the same cell type. We demonstrate the power of same-sample and same-cell multiomics analyses to further elucidate the mechanisms underlying PSC cell type and cell state in both sexes and throughout the age span.

## Methods

### Sample procurement

Flash-frozen *post-mortem* human pituitaries were obtained from the National Institutes of Health (NIH) NeuroBioBank, and kept at −80C until processing. See **Supplementary Table 1** for information on subject sex, age, ethnicity, *post-mortem* interval (PMI), cause of death, and year of collection. The tissues received varied from whole to pieces of pituitaries. All specimens were obtained from deceased individuals. Donor anonymity was preserved, and guidelines were followed regarding consent, protection of human subjects, and donor confidentiality.

### Nuclei isolation from pituitaries

Two methods were tested for nuclei isolation. Frozen *post-mortem* human pituitaries were either: 1) broken into small pieces in a frozen mortar on dry-ice, and one piece was thawed on ice and prepared for nuclei extraction based on a modified protocol from^44^, or 2) pulverized and part of the powder used for nuclei isolation. The remainder of the pituitary was stored back at −80C. Briefly, and all on ice, RNAse inhibitor (NEB cat# MO314L) was added to the homogenization buffer (0.32 M sucrose, 1 mM EDTA, 10 mM Tris-HCl, pH 7.4, 5mM CaCl_2_, 3mM Mg(Ac)_2_, 0.1% IGEPAL CA-630), 50% OptiPrep (Stock is 60% Media from Sigma; cat# D1556), 35% OptiPrep and 30% OptiPrep right before isolation. Each pituitary was homogenized in a Dounce glass homogenizer (1ml, VWR cat# 71000-514), and the homogenate filtered through a 40 mm cell strainer. An equal volume of 50% OptiPrep was added, and the gradient centrifuged (SW41 rotor at 17,792xg; 4C; 25min). Nuclei were collected from the interphase, washed, resuspended either in 1X nuclei dilution buffer for snATACseq (10X Genomics) or in 1X PBS/0.04% BSA for snRNAseq, and counted (Cellometer).

### SnRNAseq assay

SnRNAseq was performed following the Single Cell 3’ Reagents Kits V3 User Guidelines (10x Genomics, Pleasanton, CA). Nuclei were filtered and counted on a Countess instrument. A minimum of 1,000 nuclei were targeted (Chromium Single Cell 3’ Chip kit A v2 PN-12036 or v3 chip kit B PN-2000060). Reverse-transcription (RT) was performed in the emulsion, cDNA amplified, and libraries constructed with v3 chemistry. Libraries were indexed for multiplexing (Chromium i7 Multiplex kit PN-12062).

### SnRNAseq data analysis

SnRNAseq data were processed using the Cell Ranger pipeline v5.0.0, and aligned to the Cell Ranger GRCh38 reference genome, introns included. Clustering and differential gene expression analysis were performed using Seurat v.3.9.9.9024 and standard procedures^45, 46^. Top markers for each cluster were compared to known markers of pituitary cell types to annotate the clusters; a list of the most common genes associated to each cell type is given in **Supplementary Table 4**.

We used the t-SNE projection to identify the most common cross-type doublets, as well as apoptotic and low-count cells as t-SNE preserves the local structure of the data better than the UMAP projection. Doublet clusters appear as small, high UMI count satellites to the main clusters. We verified the nature of every such group of cell barcodes by plotting their gene expression of the top cell-type markers. By looking at which two gene expression programs are expressed in the barcodes composing each one of these satellite clusters, we were able to identify the two cell types that constitute the barcodes of these sub-clusters.

Apoptotic cells form their own clusters separate from the parent cluster. Several cell types often merge into a single apoptotic cluster, so that not every cell type will have its corresponding apoptotic cluster. These cells are characterized by low UMI counts and almost exclusively spliced mRNA reads, suggesting condensation of the nuclei and arrest of transcription of new mRNA.

Some cell-type clusters have offshoots composed of barcodes with low UMI counts. Contrary to apoptotic clusters, these have a similar ratio of intronic to exonic reads as their parent cluster and do not form their own cluster, but usually connect to, or appear very close to, their parent cluster. These are probably experimental artifacts of slow mRNA capture. Their gene expression program is the same as that of their higher UMI counterparts, but they are more adversely affected by dropouts. As such, we decided to remove these low-UMI offshoots from downstream analysis, together with doublet barcodes and apoptotic cells.

### SnATACseq assay

SnATACseq was performed following the Chromium Single Cell ATAC Reagent Kits V1 User Guide (10x Genomics, Pleasanton, CA). Nuclei were counted (Countess counter), transposition was performed in 10 μl at 37C for 60min on at least 1,000 targeted nuclei, before loading of the Chromium Chip E (PN-2000121). Barcoding was performed in the emulsion (12 cycles) following the Chromium protocol. Libraries were indexed for multiplexing (Chromium i7 Sample Index N, Set A kit PN-3000262).

### SnATACseq analysis

SnATACseq data were processed using Cell Ranger-ATAC pipeline version 1.2.0, and aligned to the Cell Ranger-ATAC GRCh38 reference genome. Clustering was performed using Seurat/Signac versions 3.1.5/0.2.4 and standard procedures^47^. We produced chromatin accessibility tracks around known pituitary cell type marker genes and looked for promoter accessibility of these genes to annotate the clusters.

Doublets, low-count, and apoptotic cells were identified in the same manner as for snRNAseq data, except that for ATAC data, the UMAP projection works better and was used instead. We used the number of fragments in peaks as an indicator as to whether a cell was healthy or a doublet or low-count / apoptotic. Doublets were checked to possess fragments in peaks associated to the main markers of both cell types. In general, we have many fewer cell barcodes in the ATAC data, so doublets are also less common. Consequentially, few doublet clusters were identified.

### Sn multiome assay

Sn multiome was performed following the Chromium Single Cell Multiome ATAC and Gene Expression Reagent Kits V1 User Guide (10x Genomics, Pleasanton, CA) on part of the pulverized pediatric female sample. Nuclei were counted (Countess counter), transposition was performed in 10 μl at 37C for 60min targeting 10,000 nuclei, before loading of the Chromium Chip J (PN-2000264) for GEM generation and barcoding. Following post-GEM cleanup, libraries were pre-amplified by PCR, after which the sample was split into three parts: one part for generating the snRNAseq library, one part for the snATACseq library, and the rest was kept at −20C. SnATAC and snRNA libraries were indexed for multiplexing (Chromium i7 Sample Index N, Set A kit PN-3000262, and Chromium i7 Sample Index TT, Set A kit PN-3000431 respectively).

### Sn multiome analysis

We analyzed the pediatric female sample by sn multiome. We pooled together libraries from both GEM wells and ran the Cell Ranger ARC 1.0.0 pipeline on the pooled sample following 10x Genomics guidelines. Running the pipeline on each GEM well separately reveals that the samples have 97 barcodes in common that are called as cells.

We used Seurat version 3.9.9.9024 with Signac version 1.1.0 to perform our clustering analysis using a weighted shared nearest neighbor graph approach. This method identifies, for each cell, its nearest neighbors based on a weighted combination of the two modalities (Gene Expression & Chromatin Accessibility). We similarly use the weighted nearest neighbor graph to obtain a UMAP projection of the data. The Gene Expression modality was used to identify cluster cell types after determination of top markers for each cluster.

Apoptotic, low-count cells and doublets were also identified in a manner analogous to that of snRNAseq data. Both apoptotic and low-count cells were identified as having much lower counts of both their number of transcripts as well as their number of fragments overlapping ATAC peaks. Apoptotic cells further have a large proportion of mitochondrial gene transcripts, whereas low-count cell transcripts are dominated by background genes. Doublets, on the other hand, were identified as having higher counts in both RNA and ATAC, and expressing gene programs of two cell types simultaneously. Only one of the main clusters was identified as doublets, and no further sub-clustering was attempted.

### Merged datasets analysis

All male and female samples were merged by sex in Seurat at the UMI count level, and all of the clustering analysis was repeated on the merged samples independently from the beginning. We followed the same analysis steps as for individual samples. Unlike our integrated samples (see later section), the merged samples do not have batch effects removed. Despite that, we do not observe any systematic batch effect between our samples. We do, however, see differences in gene expression from one subject to another among some specific cell types. The merged samples allow us to highlight these differences in the implicated cell types.

### Quality control (QC) and sequencing of libraries

Libraries were quantified by Qubit 3 fluorometer (Invitrogen) and quality was assessed by Bioanalyzer (Agilent). Equivalent molar concentrations of libraries were pooled and the reads were adjusted after sequencing the pools in a Miseq (Illumina). The libraries were then sequenced in a Novaseq 6000 (Illumina) at the New York Genome Center (NYGC) following 10X Genomics recommendations.

### Mouse snRNAseq stem cell data analysis

SnRNAseq data for each mouse were processed and analyzed as previously described^13^. Raw reads from each mouse sample were isolated from the clusters assigned in Seurat as ‘Stem cells’ using the ‘WhichCells’ function. These count tables were integrated using Seurat (v3.1.5) SCTransform workflow^48^, clustered at 0.5 resolution and principal component dimensions 1:15 were taken forward for analysis.

### Human stem cell re-clustering method

Following initial clustering of the complete datasets, the ‘stem cell’ clusters were isolated from each individual subject using the Seurat ‘subset’ function. To increase the number of cells available for downstream analysis, the isolated stem cell datasets were merged based on the approximate age of subjects. This was performed using the merge function within Seurat (v3.1.5). Sample integration by identification of anchors and subsequent clustering (20 PCAs, resolution 0.5) was performed using Seurat according to standard procedures^45, 46^. Re-clustering and analysis leading to identification of ‘committing’ stem cells was done as above following removal of the ‘Pars Tuberalis’ cell clusters.

### RNAscope mRNA *in situ* hybridization

Wildtype CD-1 murine postnatal pituitaries were dissected at P3, P15 (male), and P56 (male), and fixed in 10% neutral buffered formalin (Sigma) at room temperature for 16-24 hours. Samples were washed in PBS and dehydrated through graded ethanol series before paraffin-embedding as previously described^15^. Samples were sectioned at 5 μm.

The RNAscope 2.5 HD Duplex assay (Advanced Cell Diagnostics) was used according to manufacturer’s recommendations, with the following specific probes: Mm-Jun (Cat# 453561) and Mm-Sox2-C2 (Cat# 401041-C2) (Advanced Cell Diagnostics). Sections were counterstained with Mayer’s hematoxylin (Vector H-3404) and mounted with Vectamount Permanent Mounting Medium (Vector H-5000).

### Pseudotime sn trajectory analysis

Raw gene counts were extracted from each sample’s “Stem Cell” cluster as previously identified using Seurat (v4.0.1)^49^. Due to sex differences, male and female samples were handled separately. Samples of the same sex (e.g female pediatric, female adult, female aged), were integrated using the functions ‘*SCTransform*’ and ‘*SelectIntegrationFeatures*’ in Seurat to obtain the top 500 differentially expressed genes (DEGs). Monocle3 (v0.2.3.0)^50^ was used for pseudotime trajectory analysis and preliminary analysis revealed a bias in the Monocle trajectory due to specific hormonal genes namely; *GH1*, *PRL*, *CHGB*, *POMC*, *LHB*, *FSHB* and *CGA*. Therefore, these genes were regressed out of the top 500 DEGs. Monocle objects were generated by combining the 3 samples of each sex using the respective top 500 DEGs. The trajectory was calculated by merging partitions with the root chosen based on the earliest timepoint available for each sex. To find gene modules changing over pseudotime, the ‘*graph test*’ function was carried out using the neighbor_graph = “principal_graph” parameter with a resolution of 0.8 for ‘*find_gene_modules*’ function. The top 4 enriched modules for each age in each sex were highlighted and examined further because they showed the highest variability between age groups.

### PLIER data analysis

To examine more deeply the trends in gene expression of assigned cell types across samples and data types, we treated each sc dataset as a collection of bulk datasets for given labeled cell types. Each cell type was then treated as a separate bulk measurement within each sample. For snATACseq data, peak counts for a given gene were generated by selecting the peak closest to the transcription start site (TSS). These peak counts per gene were then collected into single bulk measurements for each cell type in each sample. We focused specifically on six relevant cell types in the pituitary: corticotropes, gonadotropes, lactotropes, somatotropes, stem/progenitor cells, and thyrotropes. For the snRNAseq dataset, this process generated 36 bulk measurements over six samples (three females and three males), and for the snATACseq dataset, we generated 35 bulk measurements as thyrotropes were not identified in the male adult snATACseq sample. We applied PLIER^29^, which finds patterns in count data that are associated with known prior information (such as Reactome and Kegg), focusing on the 2000 genes with the highest standard deviation in count values across the bulk measurements in each set of samples. PLIER was run on each set of samples separately with LVs generated on the bulk measurements in an unsupervised fashion. LVs were then curated to find patterns relevant to individual cell types as well as sample-wide trends such as sex-based differences. Statistical significance of LVs was computed through the Kruskal-Wallis non-parametric test for multiple groups as part of the stat_compare_means R method. Comparisons between LVs within and across datatypes were achieved by comparing the overlap of the 200 genes most associated with a given LV.

B is a PLIER-derived expression value for the genes associated with a given LV across the different samples. It can be treated similar to average expression, weighted by gene association with the LV. Technically, B is a matrix of size #LVs x #Samples. It is one of two matrices in PLIER, along with Z of size # of genes x #LVs. The goal of PLIER is to find values of B and Z that minimize the equation ||Y - Z*B|| where Y is our data matrix of size #genes x #samples. So PLIER finds a suitable number of LVs that can be used to connect the genes and samples and accurately estimate our data matrix.

For the boxplot statistical analysis (**Supplementary Fig.12**), ggboxplot generates a boxplot with the center equal to the 50th percentile, the bounds of the box are the 25th and 75th percentile and the bounds of the whiskers are the smallest/largest values 1.5 times the interquartile range below the 25th percentile or above the 75th percentile, respectively.

### Sn data integration

The snRNAseq and snATACseq data were integrated in a reference-query based manner, mainly using the “FindTransferAnchors” and “TransferData” functions from the Seurat v3 package^45, 46^. The snRNAseq datasets were used as the reference and the other modalities were integrated to them. To integrate snATACseq to snRNAseq, the peak-by-cell accessibility matrix was converted to a gene-by-cell activity matrix based on the chromatin accessibility within each gene’s gene body and a 2kb upstream region, under the assumption that chromatin accessibility and gene expression were positively correlated. The variable features from the snRNAseq data were used to find the anchors and the snATACseq data in the LSI low-dimensional embedding were used to transfer the data from snRNAseq to snATACseq.

### Linear modeling analysis

The regulatory model of gene expression from the sn multiome dataset was constructed for each target gene with multiple steps: 1) A list of potential regulatory genomic regions were selected. They included: a) any ATAC peaks that overlaped with the TSS +/-2kb region, b) distal peaks that were no more than 500kb away from the TSS and were co-accessible with any of the peaks in a). Co-accessibility scores were calculated using the Cicero package ^51^ with default parameters, and a cutoff of 0.25 for the co-accessibility scores were used to select co-accessible peaks. 2) A list of potential regulatory TFs was selected by scanning for TF binding sites in the selected genomic regions using the “matchMotifs” function (with a p value cutoff of 5e-5) from the r package “motifmatchr” and the position weight matrices (PWMs) from the JASPAR CORE database. 3) Linear regression was used to model the target gene’s expression across cells as a function of selected TFs’ expression and ATAC peaks’ openness, and the coefficients from the regression were used to measure the importance of each TF and genomic region. SCTranform^48^ normalized RNA counts and TFIDF normalized ATAC peak counts were used in the regression.

### Statistics

In **Fig. 5a**, to calculate the statistical significance of expression or accessibility changes within a given latent variable, we applied two-way ANOVA for multiple group testing and Tukey test for pairwise comparisons. Each test was applied to female and male samples separately. In both cases, we applied the R statistical functions *aov* and *TukeyHSD* with the additive model *Expression ~ ‘Cell Type’ + ‘Age Group*’ for the calculations.

In **Supplementary Figs. 11a** and **12a**, the hierarchical clustering of the LV B scores was accomplished through the default complete linkage method utilized by the R function *pheatmap*.

For the boxplots analysis in **Supplementary Fig. 12c**, the analysis was done with n=3 independent subjects per sex and statistical analysis using the Wilcoxon ranked-sum test.

**For Fig. 6 and Supplementary Figs. 15-19**, the P-values of peaks and the P-values of the TFs were both obtained by running a linear regression (“lm” function in R) on 9,151 cells (for pan-pituitary results) and 1,623 cells (for stem cell specific results). In addition, for the TFs statistical analysis, the TFs are presented only if their Bonferroni-corrected P-values < 0.05. Detailed statistics (such as t values of linear regression) are provided in **Supplementary Table 8**.

## Data availability

The datasets (snRNAseq, snATACseq, sn multiome) generated in the present study are deposited in GEO (accession # GSE178454). The sn human pituitary multi-omics atlas can be browsed via a web-based portal accessible at snpituitaryatlas.princeton.edu. All datasets will also be deposited with the Human Cell Atlas.

## Code availability

Any computational code used in the paper is available upon request.

## Acknowledgements

This work was supported by funding from the National Institute of Health (NIH) Grant DK46943 (SCS), Medical Research Council (MRC) Grant MR/T012153/1 (CLA), and Canadian Institutes of Health Research Project Grants PJT-162343 and −169184 (DJB). TLW was funded by King’s College London as part of the “Cell Therapies and Regenerative Medicine” Four-Year Welcome Trust PhD Training Programme. We acknowledge the New York Genome Center for sequencing. Human tissue was obtained from the NIH NeuroBioBank. This work was supported in part through the computational and data resources and staff expertise provided by Scientific Computing at the Icahn School of Medicine at Mount Sinai.

## Author contributions

ZZ, MZ, GRS, VY, and OGT contributed analytic tools and analyzed data; TLW contributed analytic tools, analyzed data, and performed research; NM, VDN, NS, MAA, and MV performed research; HP drafted the manuscript; SCS designed the study, analyzed, interpreted data, and drafted the manuscript, CLA, JLT and DJB analyzed, interpreted data, and drafted the manuscript; FRZ designed the study, performed research, analyzed and interpreted data, drafted the manuscript. All authors edited the manuscript and approved its final version.

