## Supplementary Tables for "Single nucleus pituitary transcriptomic and epigenetic landscape reveals human stem cell heterogeneity with diverse regulatory mechanisms"

The Supplementary Information includes 8 Supplementary Tables.

**Supplementary Table 1: Subject characteristics**

PMI = Post-mortem interval

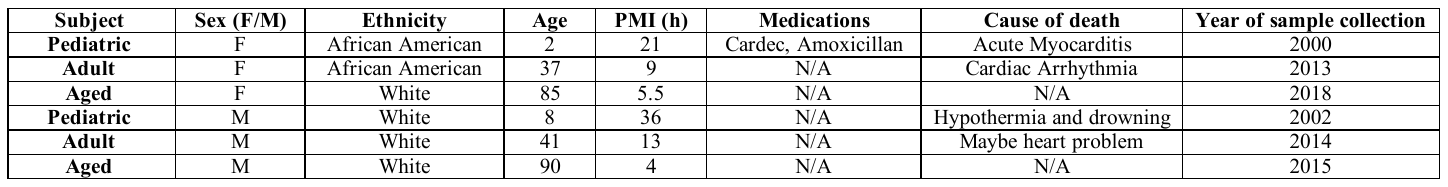

**Supplementary Table 2: Metrics obtained for the paired sn assays performed in the study.**

**a. snRNAseq metrics**

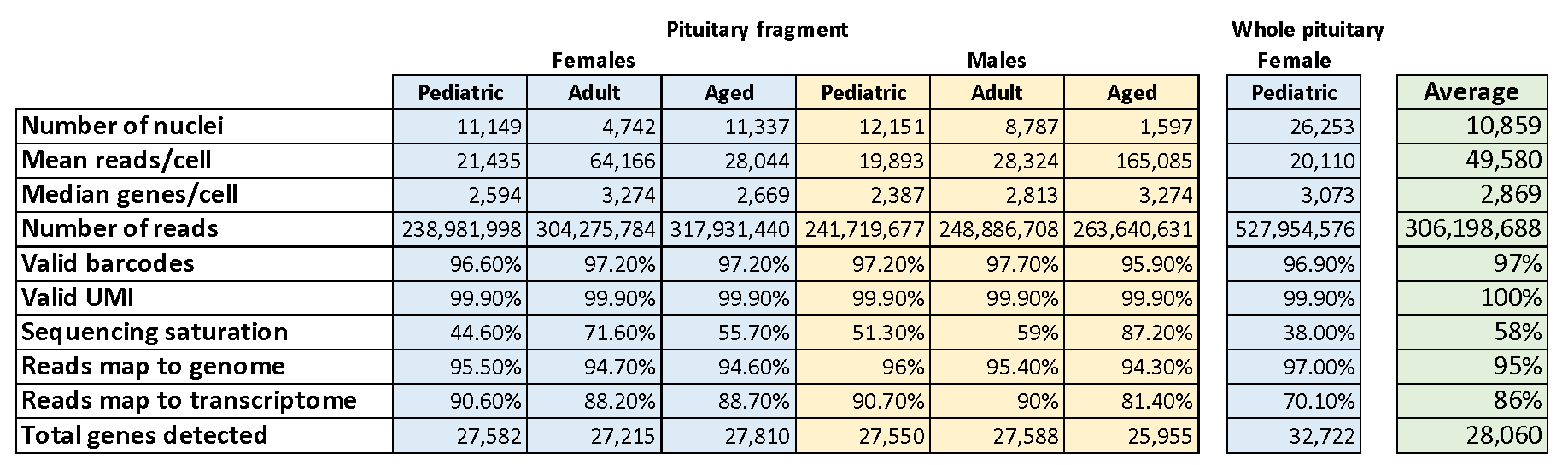

**b. snATACseq metrics**

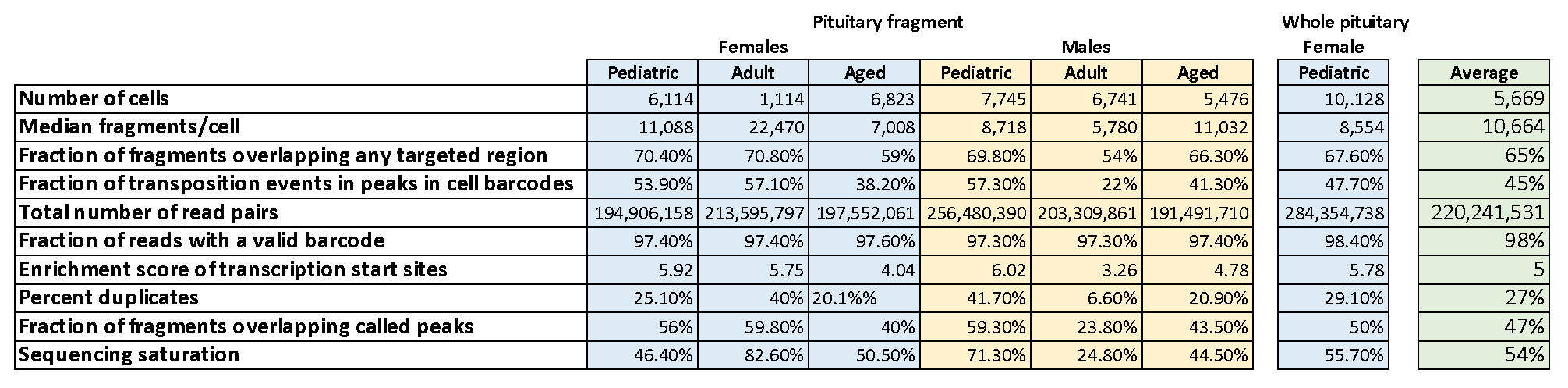

**Supplementary Table 3: Same-cell sn multiome metrics.**

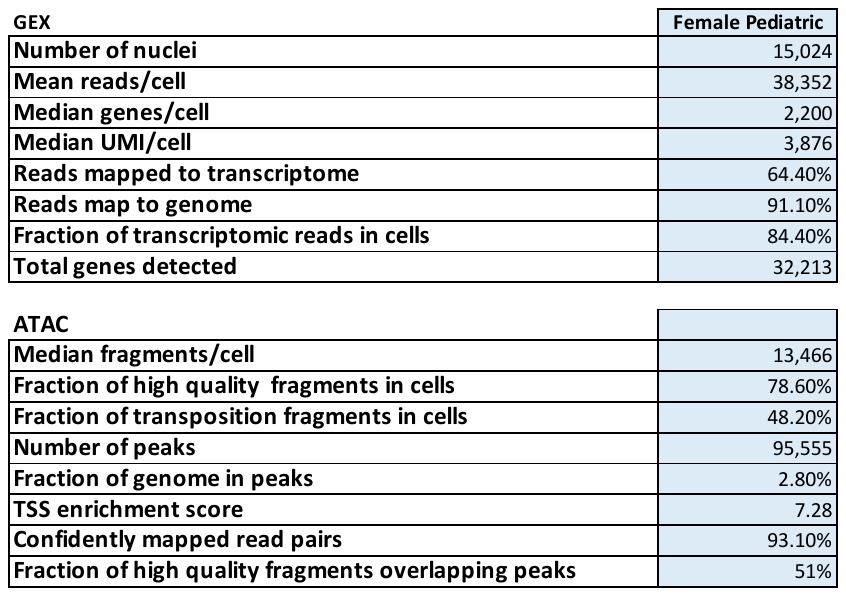

**Supplementary Table 4: Markers used for cell type assignment.**

Known cell type markers are in black. Newly identified markers from our data are in blue. Note that TSHB was only detected in a few cells in the thyrotrope cluster.

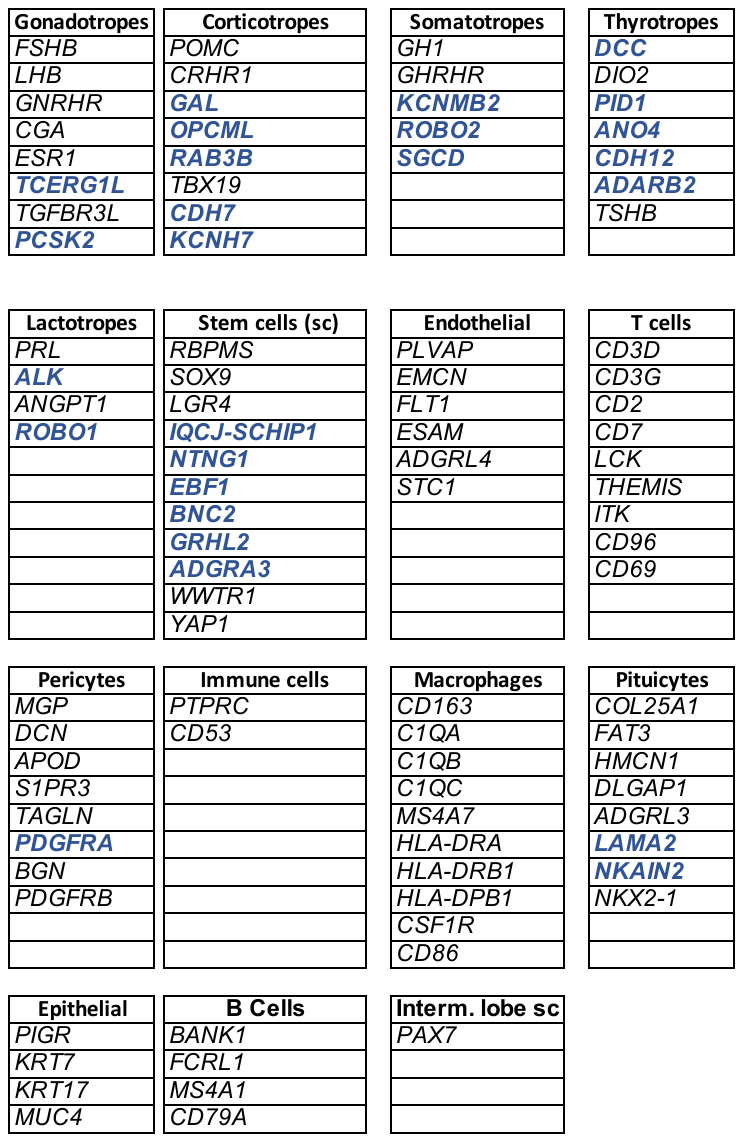

**Supplementary Table 5: Comparison of cell type proportions in the different assays done on the same pituitary sample.**

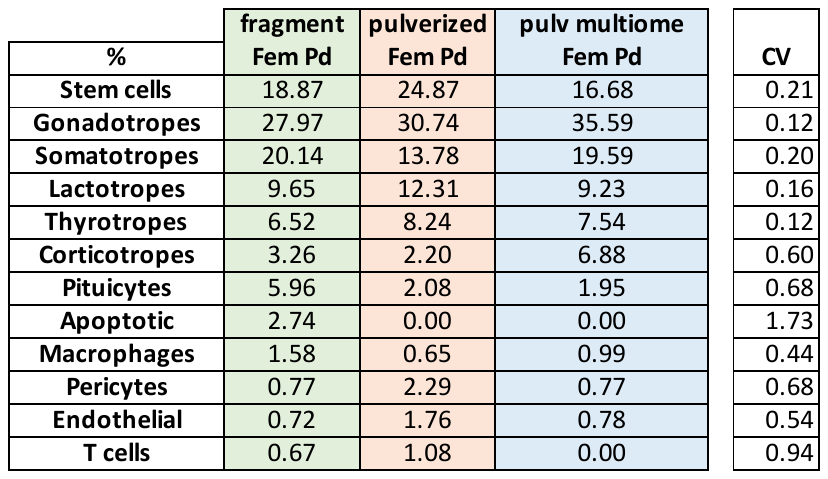

**Supplementary Table 6: Top genes per module in females (*Top*) and males (*Bottom*) for each age.**

The top 4 transcripts for each gene module are shown plus any other interesting genes present in the module. Transcripts shared across sexes highlighted in blue and transcription factors are highlighted in bold.

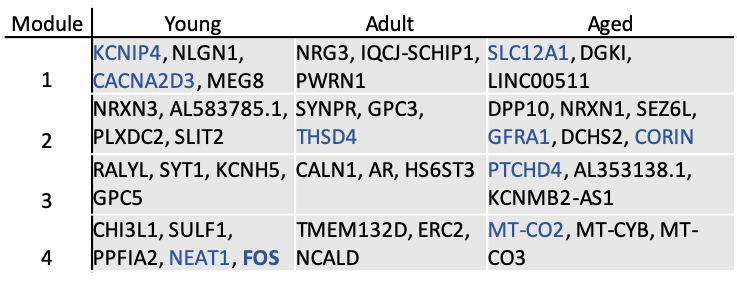

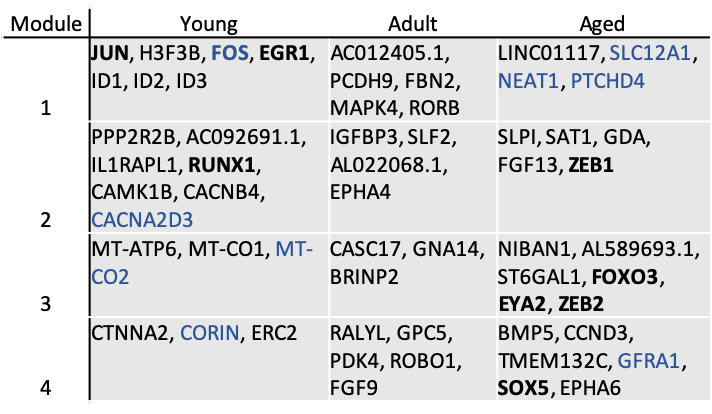

Female

Aged

Male

**Supplementary Table 7: Stem cells analyzed in snRNAseq.**

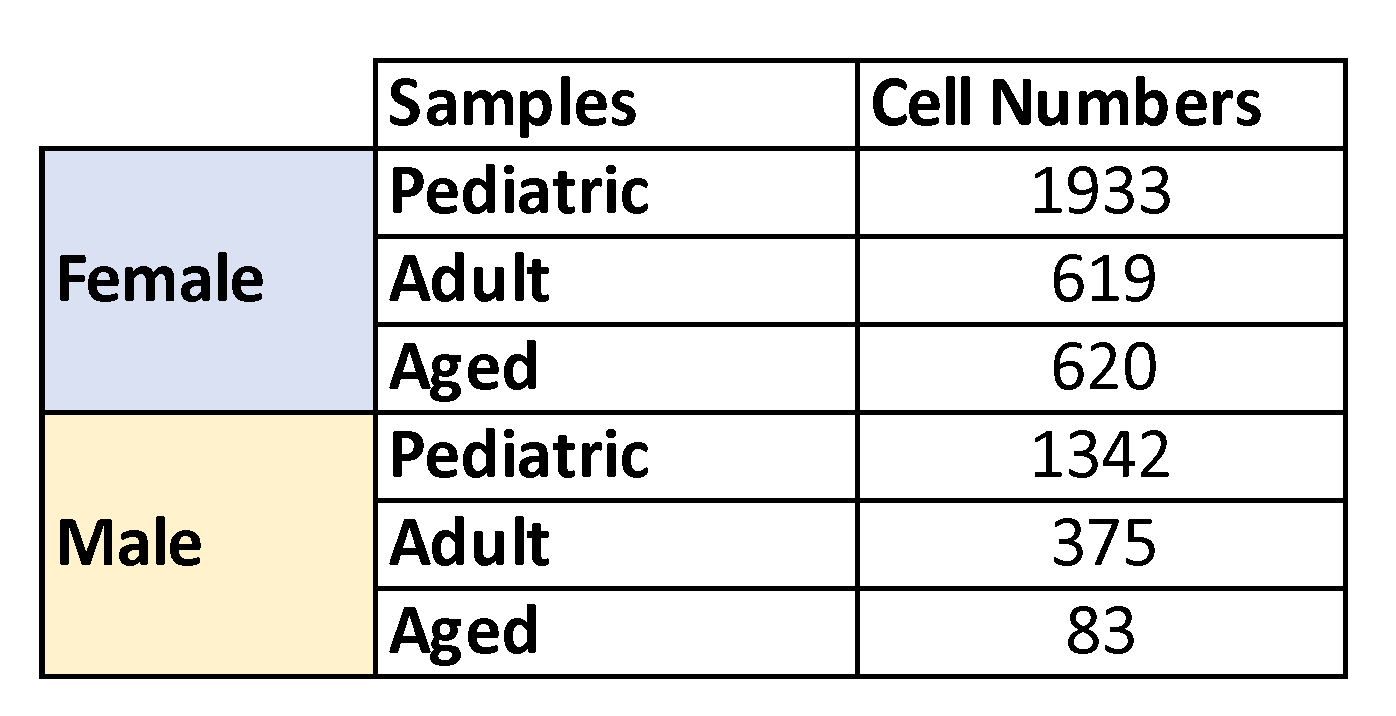

**Supplementary Table 8: Detailed statistics for the linear model analysis.**

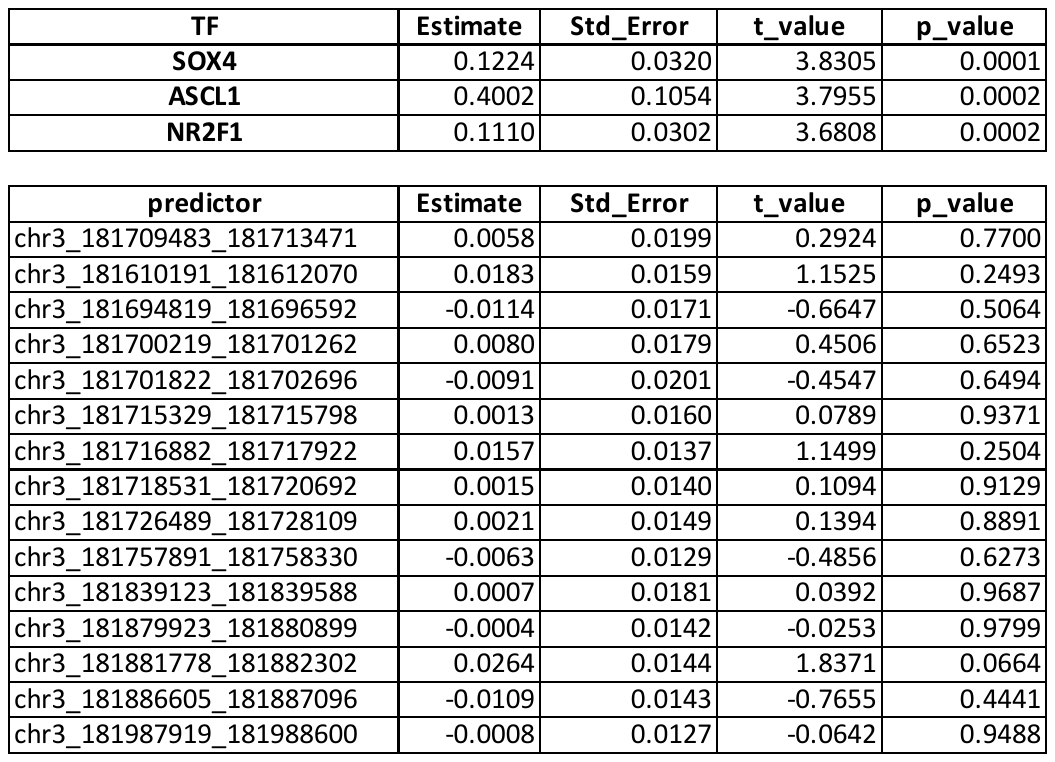
