## Supplementary Figures for "Single nucleus pituitary transcriptomic and epigenetic landscape reveals human stem cell heterogeneity with diverse regulatory mechanisms"

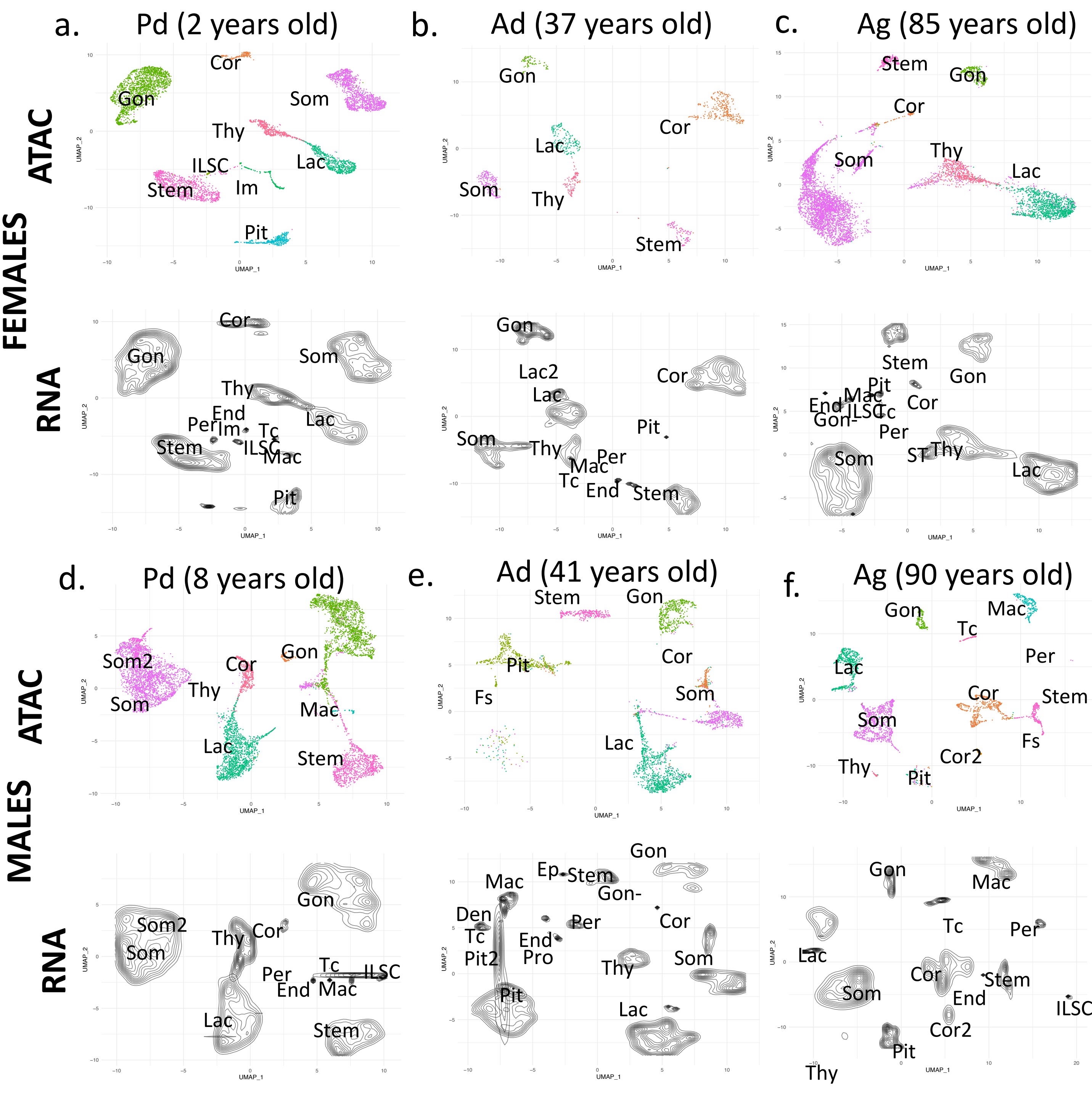


- Continuous endothelial (Cont end)
- Corticotropes (Cor)
- Corticotropes 2 (Cor2)
- Dendritic cells (Den)
- Endothelial cells (End)
- Epithelial cells (Ep)
- Intermediate lobe stem cells (ILSC)
- Gonadotropes (Gon)
- Gonadotropes - (Gon -)
- Immune cells (Im)
- Lactotropes (Lac)
- Lactotropes + (Lac+)
- Macrophages (Mac)
- Pericytes (Per)
- Pituicytes (Pit)
- Pituicytes 2 (Pit2)
- Proliferating (Pro)
- Stem cells 2 (SC2)
- Somato-thyrotropes (ST)
- Somatotropes (Som)
- Somatotropes 2 (Som2)
- Stem cells (Stem)
- T cells (TC)
- Thyrotropes (Thy)

**Supplementary Figure 1:**
Integration of the sn data modalities. For each sample, the snATACseq dataset is colored coded (top UMAP) and the snRNAseq dataset is presented as black contours (bottom UMAPs). Cell types are color-coded and designated with a letter code, as indicated on the bottom key. All integrated samples are presented in **Supplementary Figures 2**. Females (top panels; males (bottom panels), with a,d: pediatric (Pd), b,e: adult (Ad), c,f: aged (Ag)).

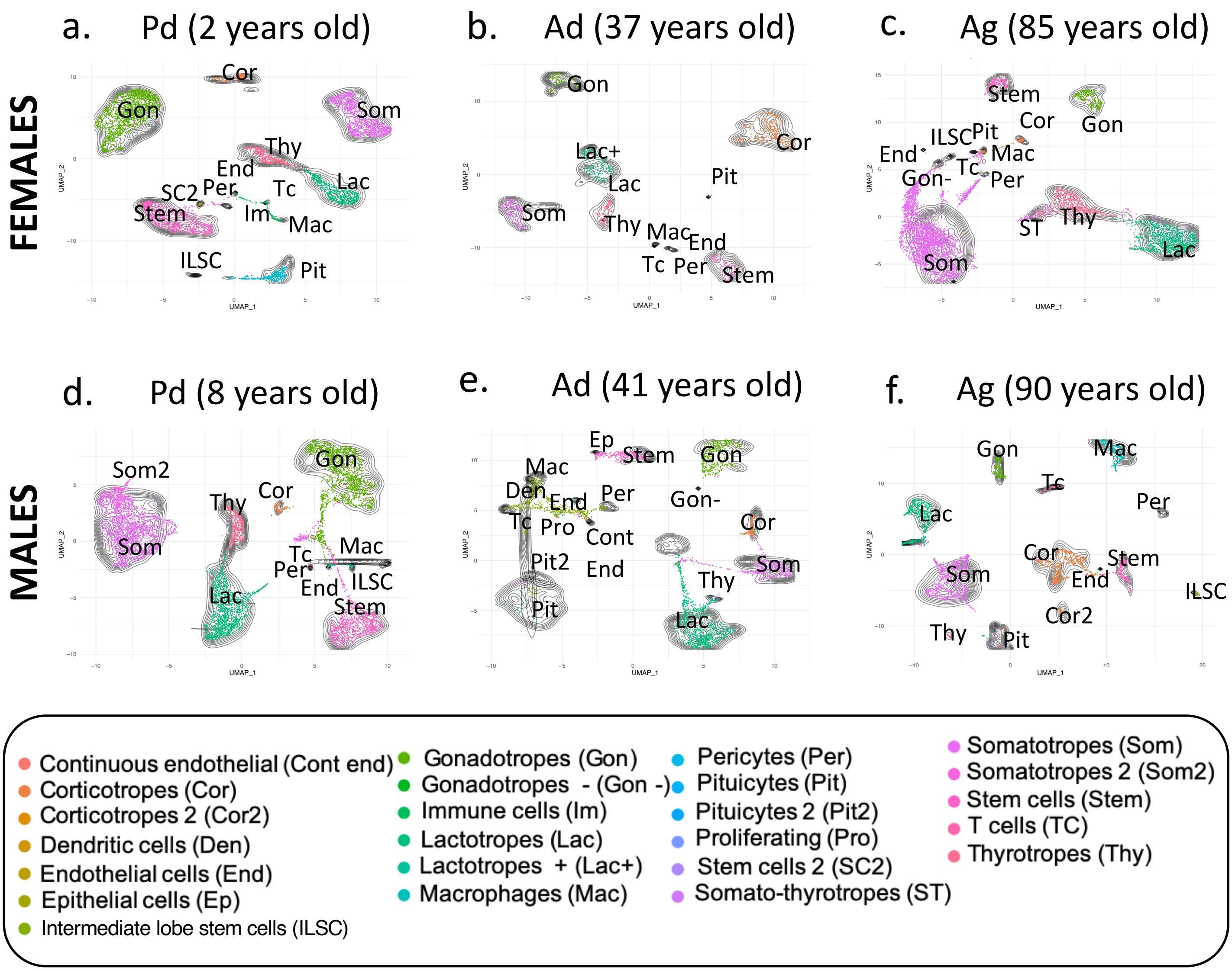


**Supplementary Figure 2:**

a-f. Integration of the sn data modalities. For each sample, the snATACseq dataset is colored coded and the snRNAseq dataset is presented as black contours on the same UMAPs. On the UMAP, cell types are color-coded and designated with a letter code, as indicated on the bottom key. Individual modalities are presented in **Supplementary Figure 1**. Females (Top panels; males (Bottom panels), with a,d: pediatric (Pd), b,e: adult (Ad), c,f: aged (Ag)).

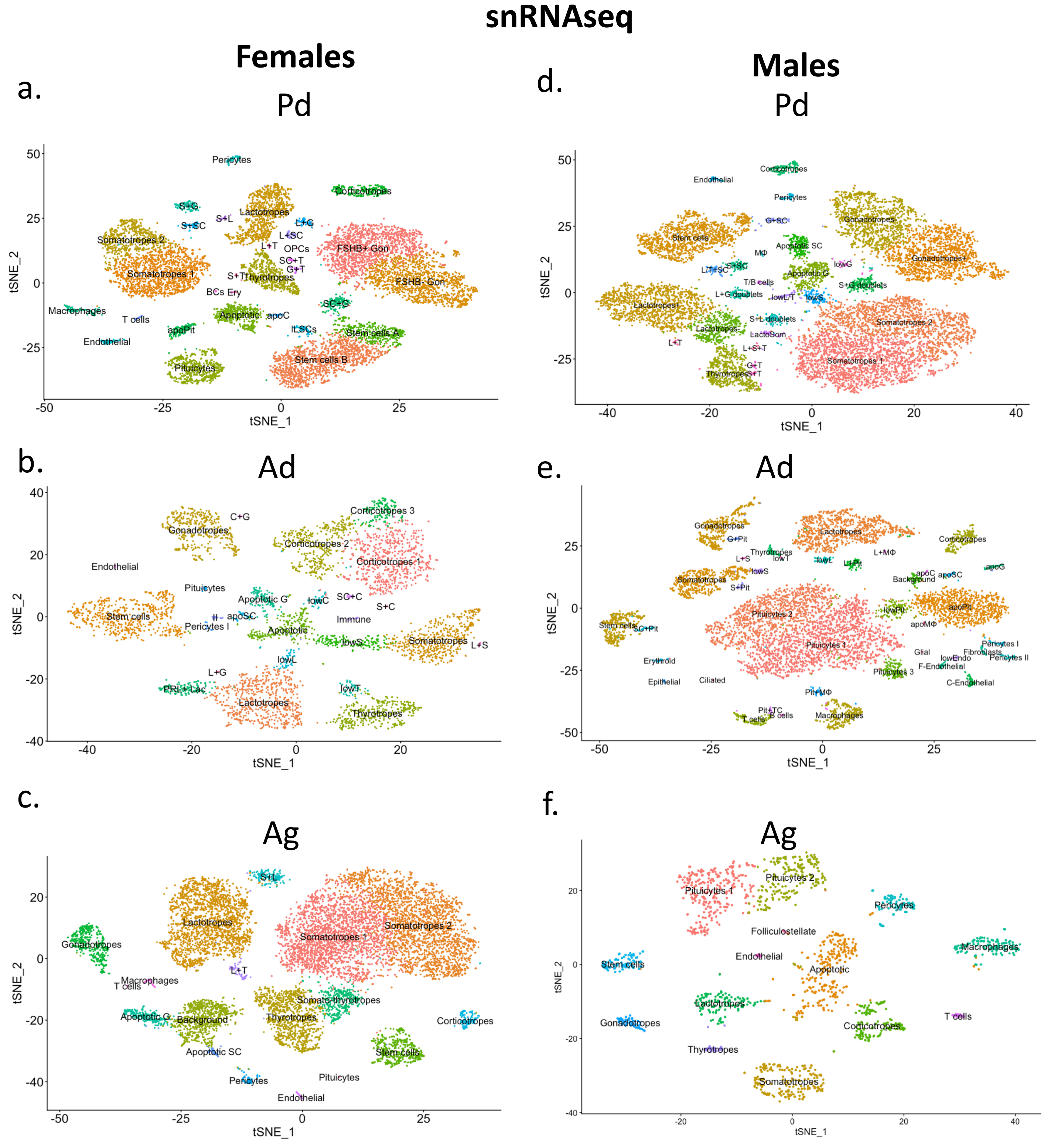


**Supplementary Figure 3:**
Results obtained for the frozen human pituitaries processed for snRNAseq. t-SNE for all individual subjects processed for snRNAseq from individually frozen pituitaries (3 females, panel a-c; and 3 males, panel d-f). a,d: pediatric (Pd); b,e: adult (Ad); c,f: aged (Ag). Refer to **Supplementary Figures 4-6** for the UMI, mitochondrial gene, and ribosomal content level analysis for all samples.

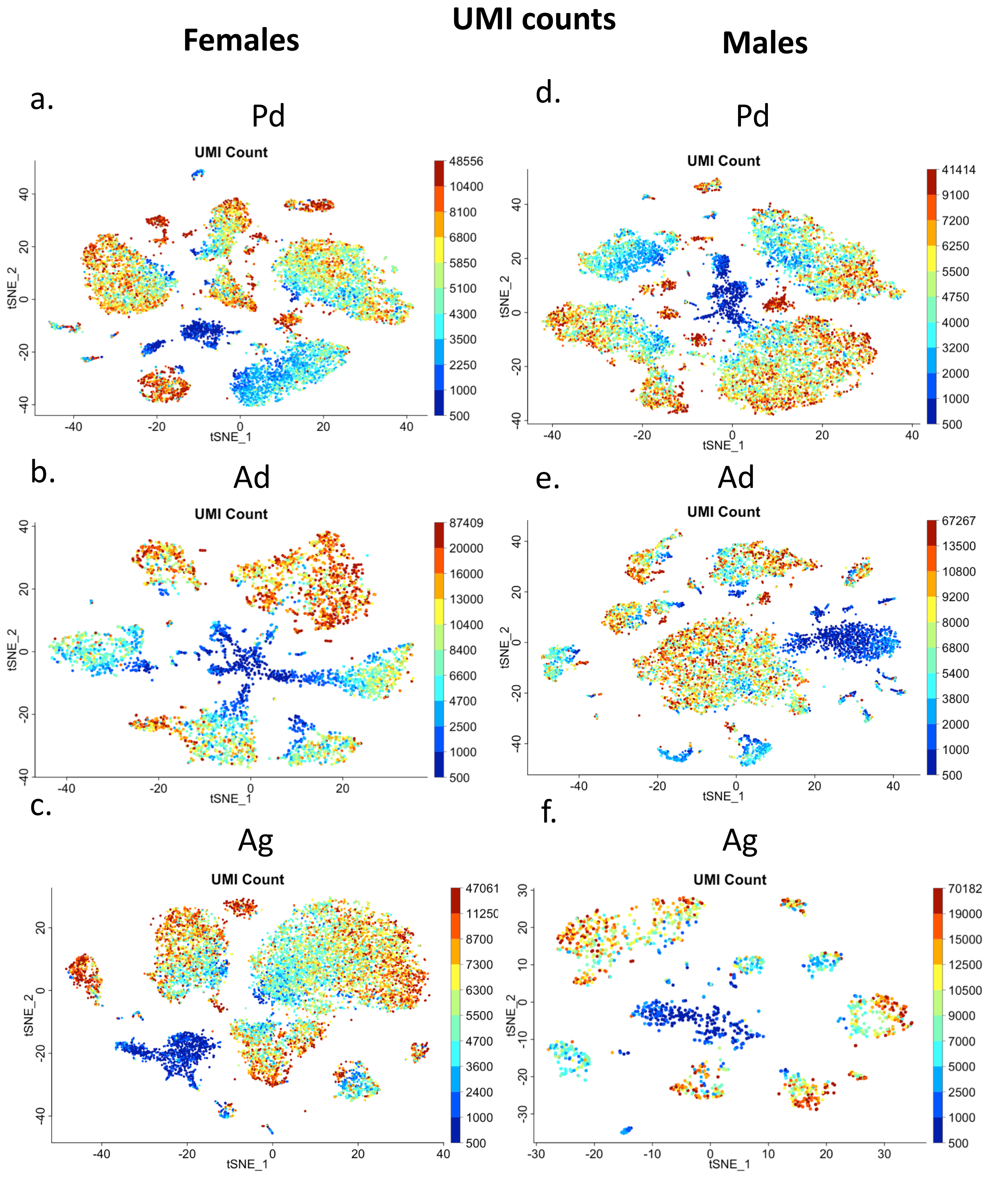


**Supplementary Figure 4: UMI count for the individual frozen human pituitary samples**

UMI counts are shown for each individual female (a-c) and male (d-f) samples after removal of doublets and debris. a,d: pediatric (Pd); b,e: adult (Ad); c,f: aged (Ag).

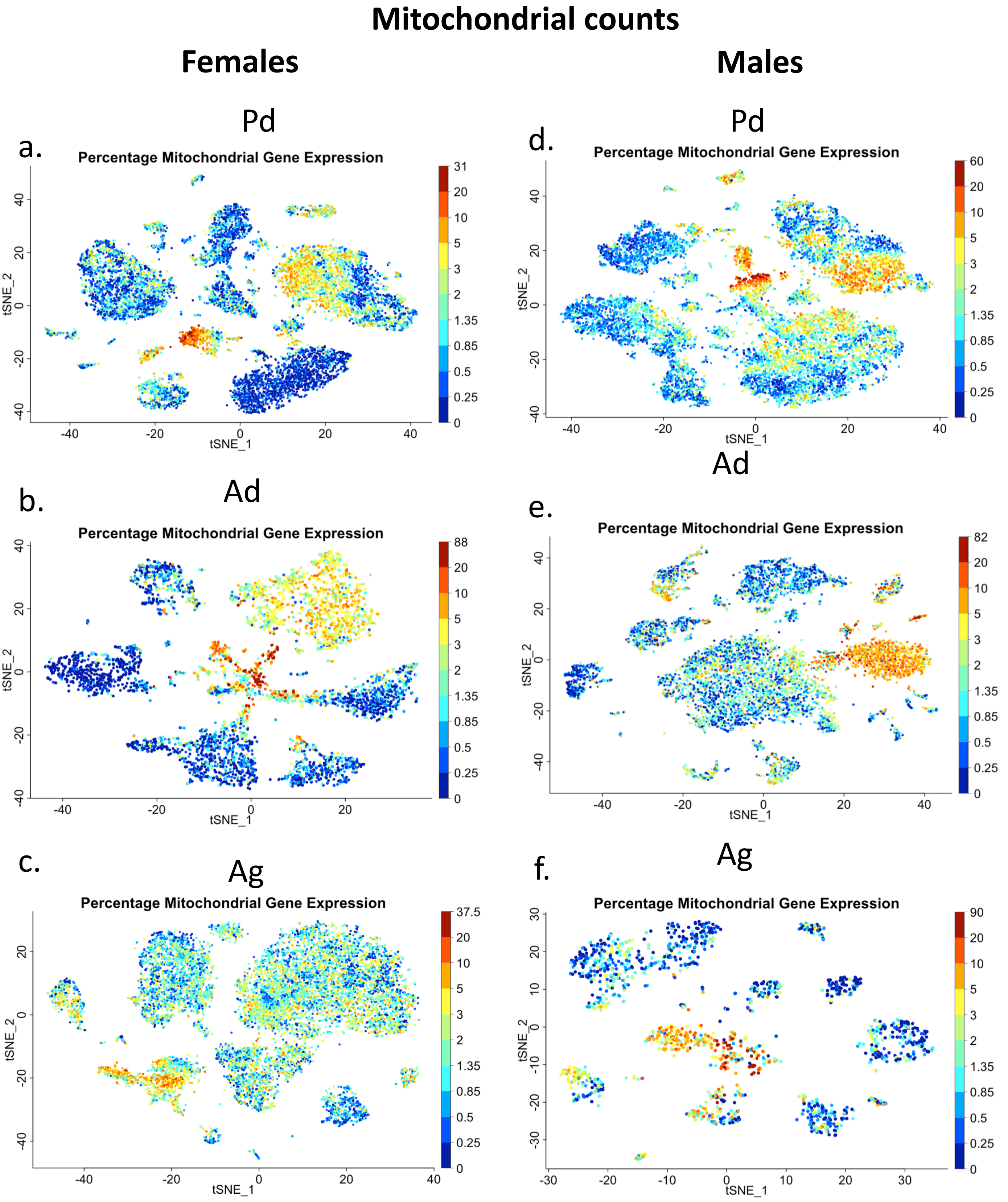


**Supplementary Figure 5: Mitochondrial content for the individual frozen human pituitary samples**
Mitochondrial gene counts are shown for each individual female (a-c) and male (d-f) samples after removal of doublets and debris. a,d: pediatric (Pd); b,e: adult (Ad); c,f: aged (Ag).

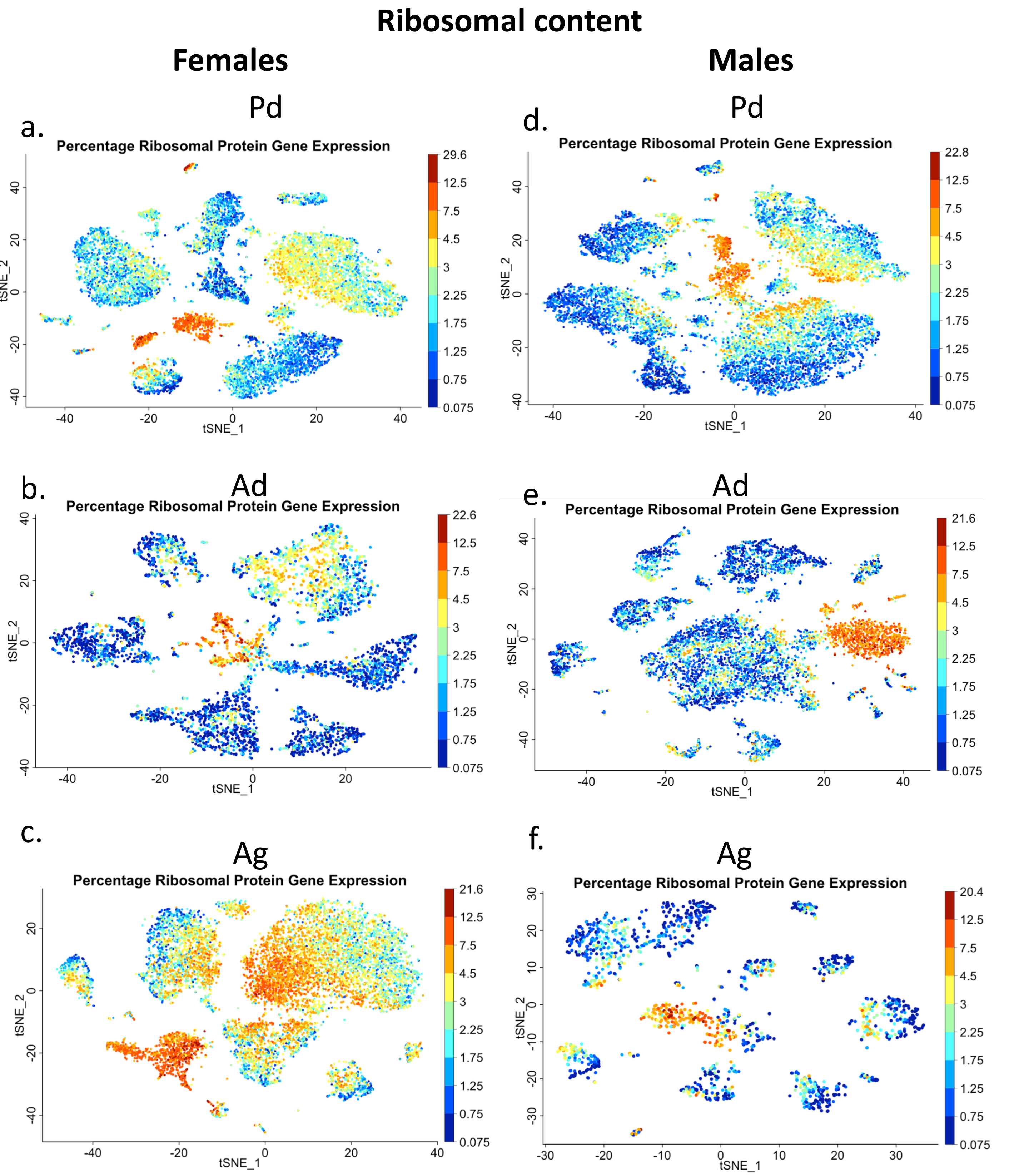


**Supplementary Figure 6: Ribosomal gene content for the individual frozen human pituitary samples**
Ribosomal gene counts are shown for each individual female (a-c) and male (d-f) samples after removal of doublets and debris. a,d: pediatric (Pd); b,e: adult (Ad); c,f: aged (Ag).

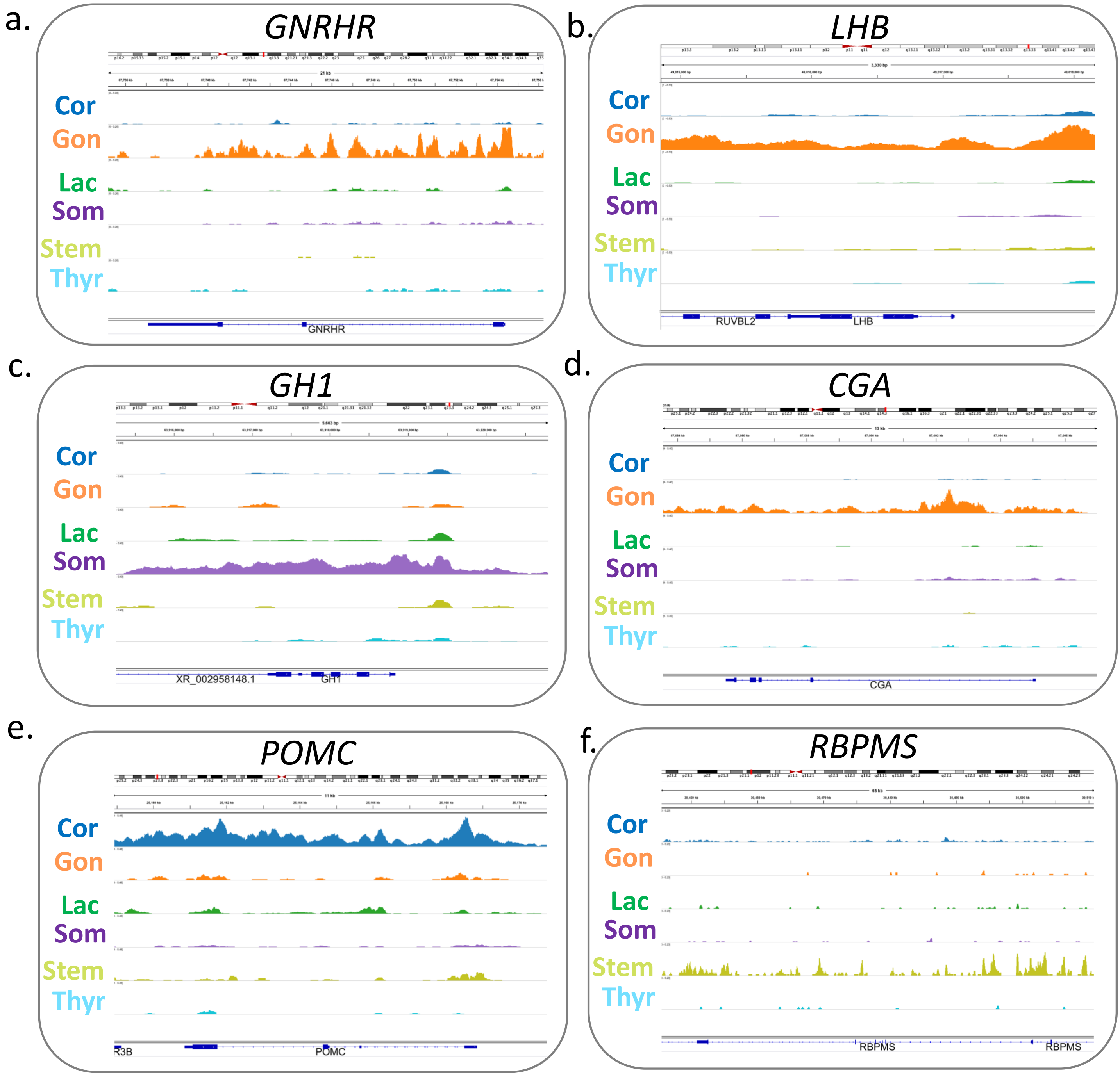


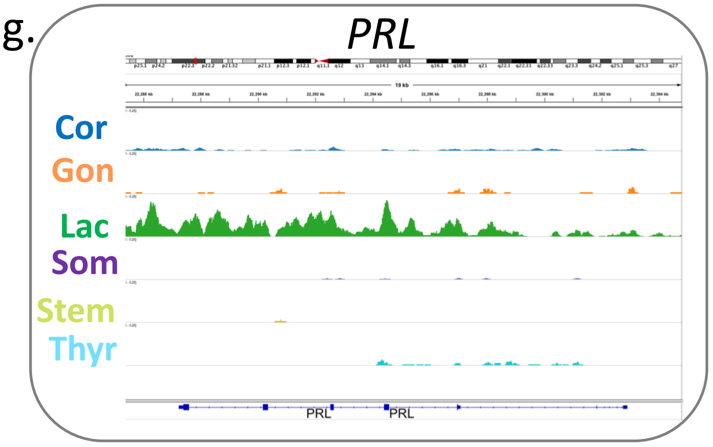


**Supplementary Figure 7:**
Shown by cell types are genome browser tracks for chromatin accessibility of selected pituitary markers. Each cell type is color coded.

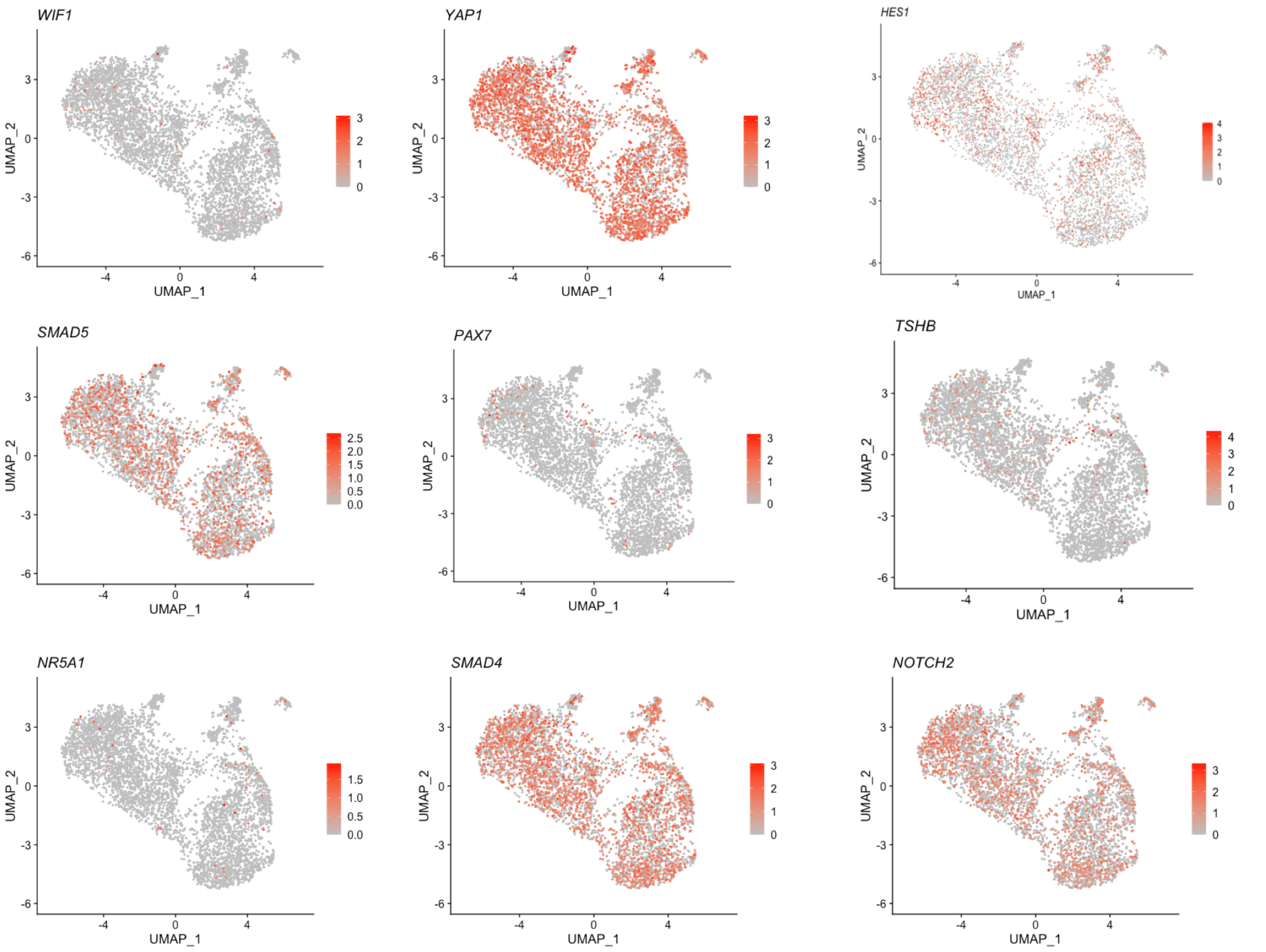


**Supplementary Figure 8:**

Feature plots of selected stem cell markers on the stem cell clusters. The intensity of the expression is represented as a red-color scale. The scale range is presented on the right of each individual plot.

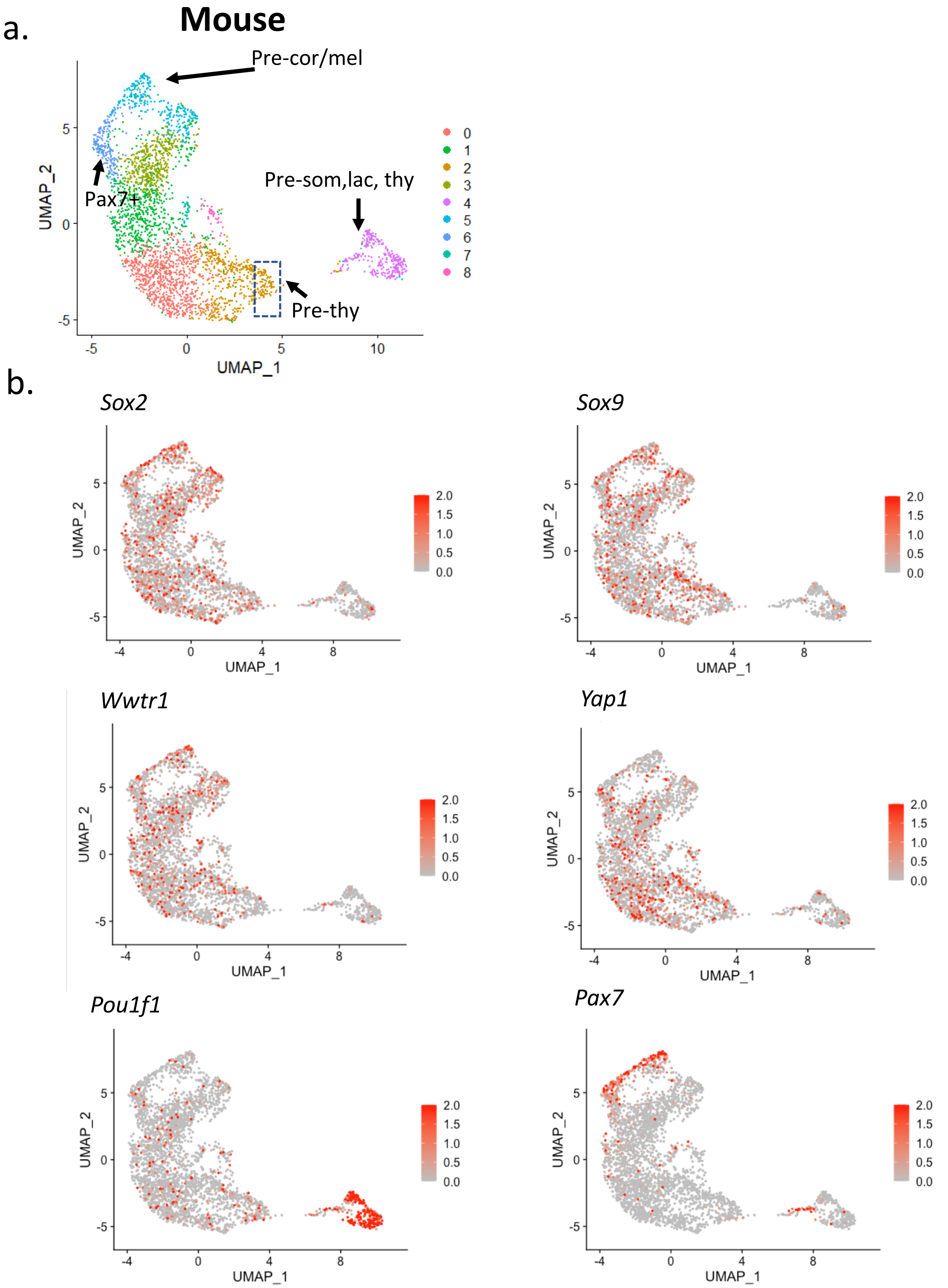


**Supplementary Figure 9:**
Analysis of the murine stem cell cluster (See Ruf-Zamojski, 2021, Nat.Comm). **a**. UMAP of the stem cell cluster identifies committing cell lineages as in human. **b.** Feature plots of selected stem cells marker expressions on the stem cell clusters. The intensity of the expression is represented as a red-color scale. The scale range is presented on the right of each individual plot.

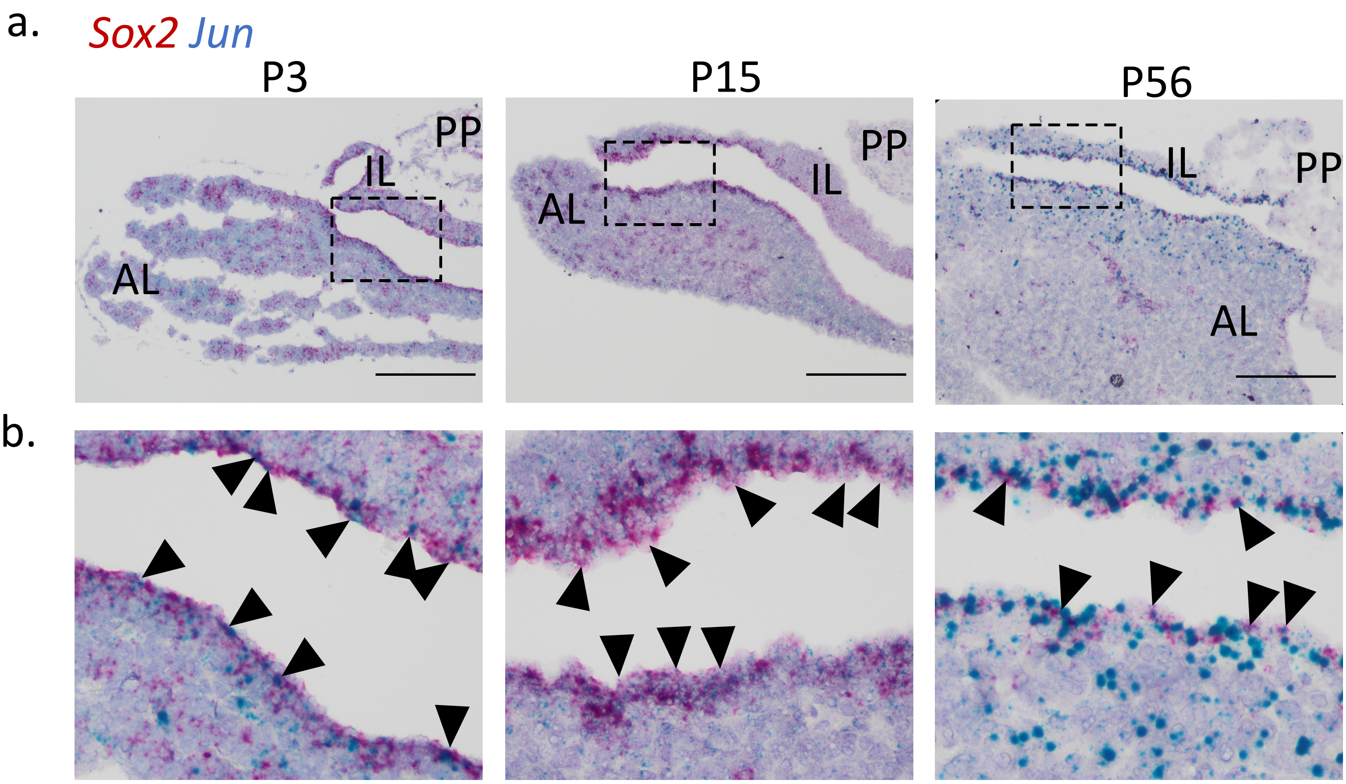


**Supplementary Figure 10:**

**a.** Wildtype murine CD-1 pituitaries show colocalization of *Sox2* (red) and *Jun* (blue) transcripts at all postnatal ages (P3, P15/male, and P56/male). Scale bar is 200µm. **b.** Boxed regions from panel **a.** are magnified. Arrows highlight specific cells with colocalization of *Sox2* and *Jun*.

AL: anterior lobe; IL: intermediate lobe; PP: posterior pituitary. The P56 panels that are included in the main figures are shown here as well for comparison.

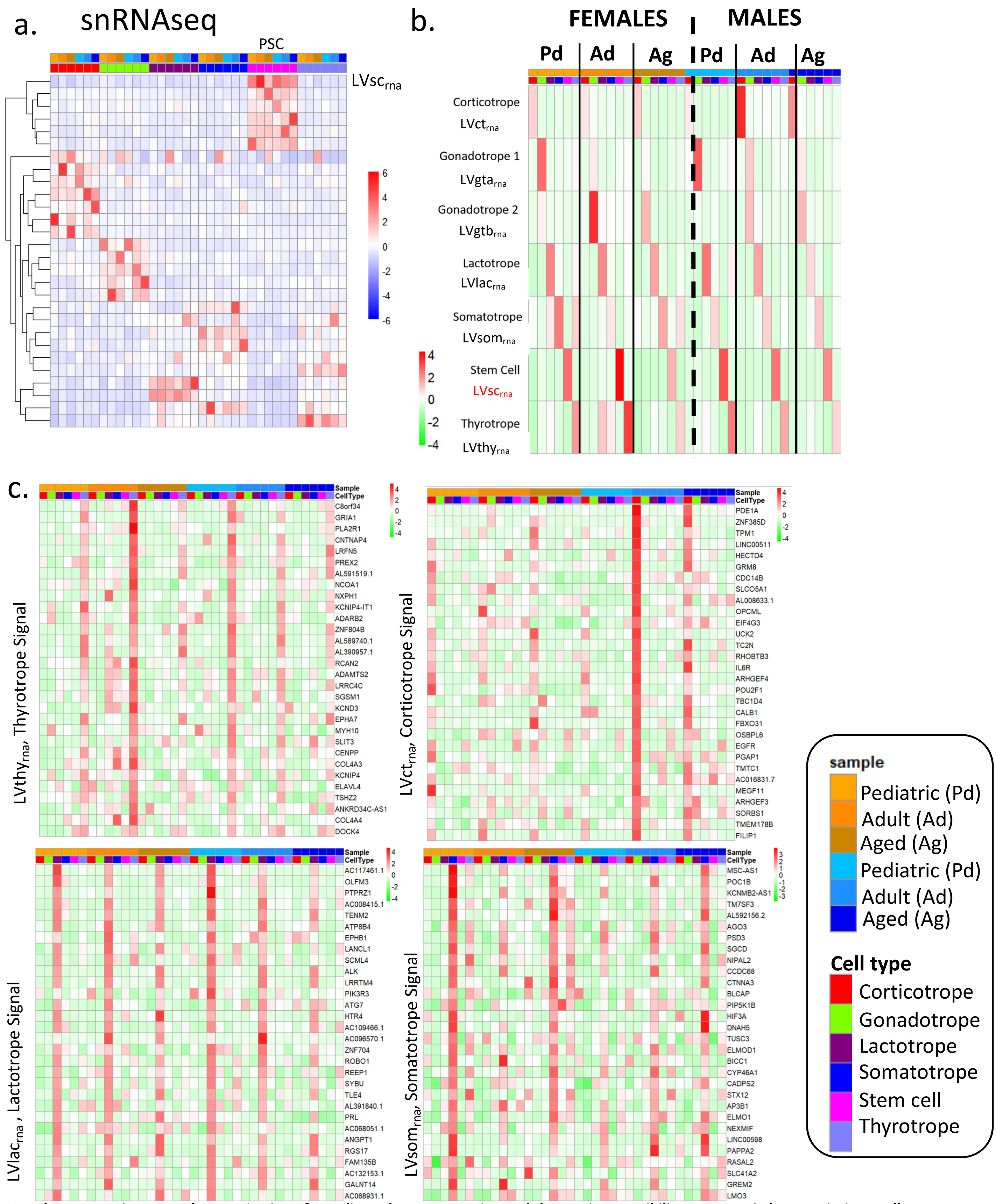


**Supplementary Figure 11: Characterization of coordinated gene expression and chromatin accessibility programs in human pituitary cell types**

**a**. Normalized data matrix of the mean raw count for each gene per cell type showing the 28 LVs identified from the 2,000 most variable genes among the subject cell types from the snRNAseq datasets. B-score values are clustered by cell type with the majority of LVs being cell type specific. **b**. Heatmaps of the levels of RNA expression of each LV for a specific cell type and/or subject. Each pituitary sample is indicated at the top. In the scale bars, red signifies the highest level of RNA expression or chromatin accessibility. Refer to Supplementary Table 1 for full information on the samples. In (**a.**) the names of the main LVs are highlighted in red. **c**. Heatmaps of the levels of gene expression for selected cell type-specific $LV_{rna}$ for each pituitary cell type and subject. The legend is presented for all heatmaps at the bottom right corner.

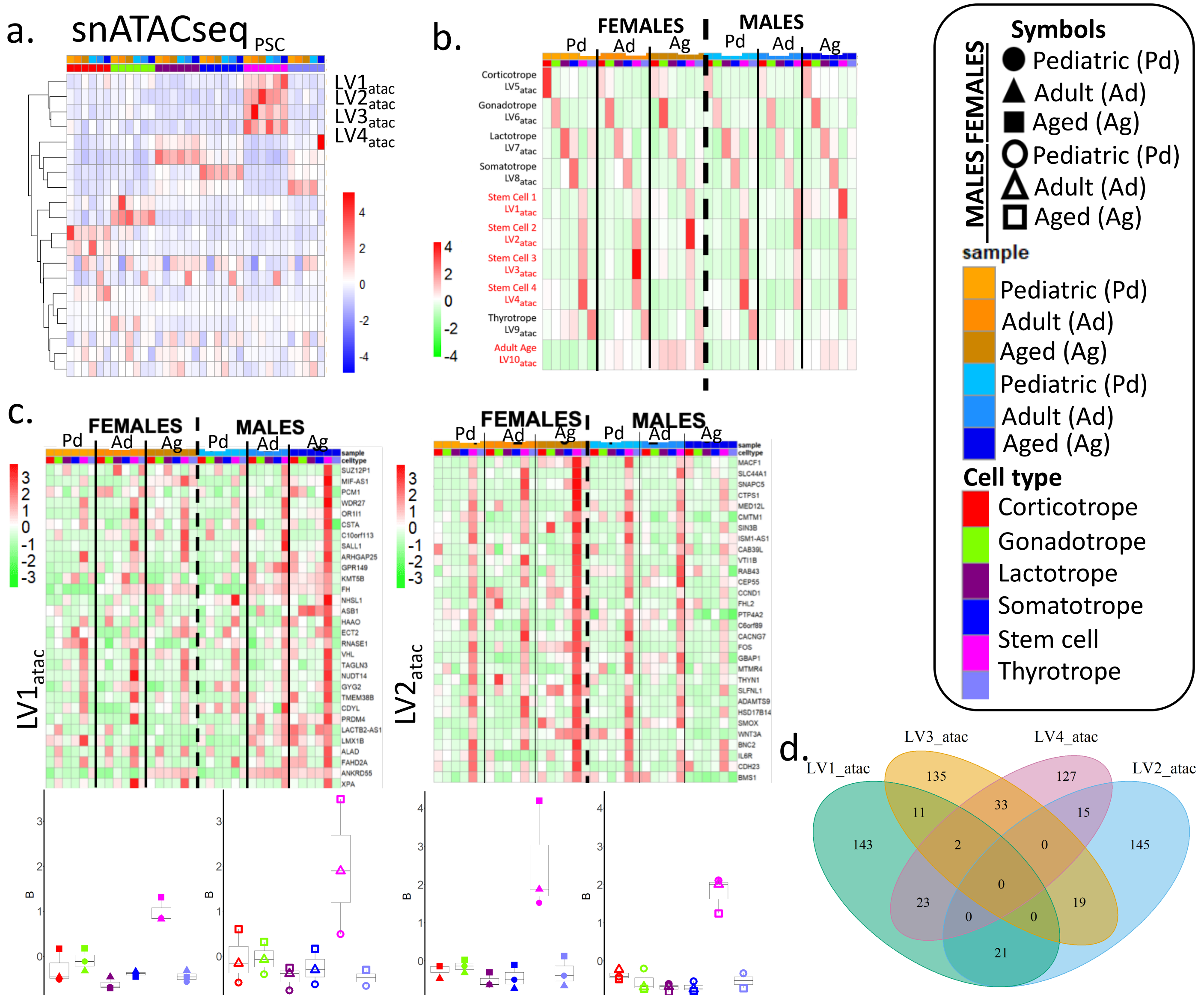


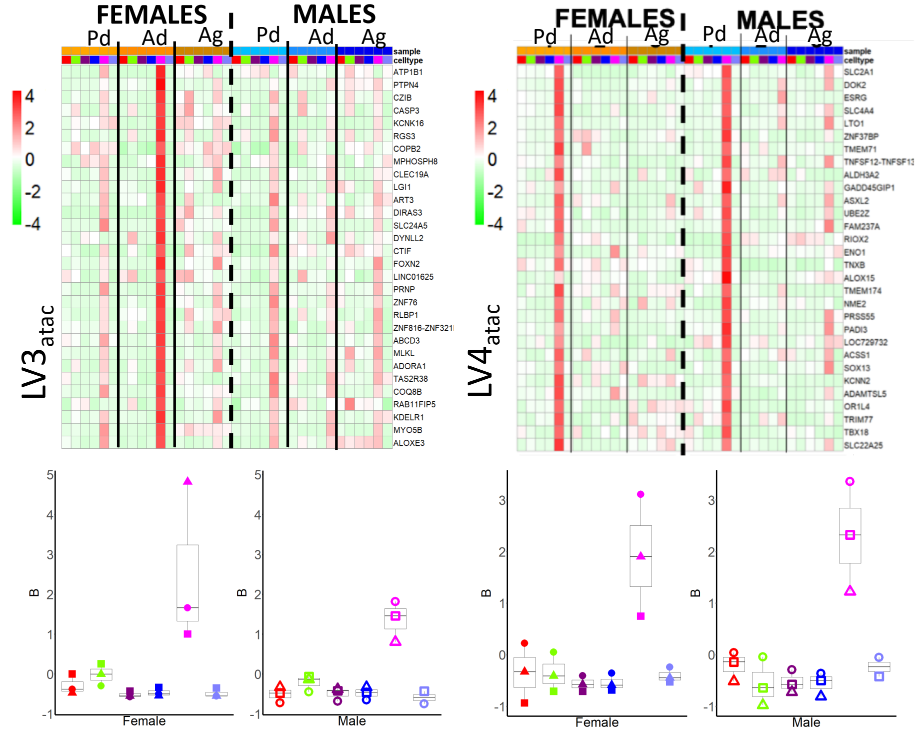


**Supplementary Figure 12: Characterization of coordinated gene expression and chromatin accessibility programs in human pituitary cell types**

**a**. Normalized data matrix of the mean raw count for each gene per cell type showing the 20 LVs identified from the 2,000 most variable genes among the subject cell types from the snATACseq datasets. B-score values are clustered by cell type with the majority of LVs being cell type specific. **b**. Heatmaps of the levels of chromatin accessibility of each LV for a specific cell type/subject. **c**. Heatmaps of the levels of chromatin accessibility for the four PSC LVs for each cell type/animal (top panels). Bottom panels are boxplots showing the statistical analysis (see Methods), with n=3 independent subjects used per sex and statistical analysis using the Wilcoxon ranked-sum test. (**a**,**b**,**c**) The full legend for symbols, cell types, and samples is presented on the right of the figure. Each pituitary sample is indicated at the top. In the scale bars, red signifies the highest level of RNA expression or chromatin accessibility. **d**. Gene overlap of the four stem cell-specific chromatin accessibility-based LVs.

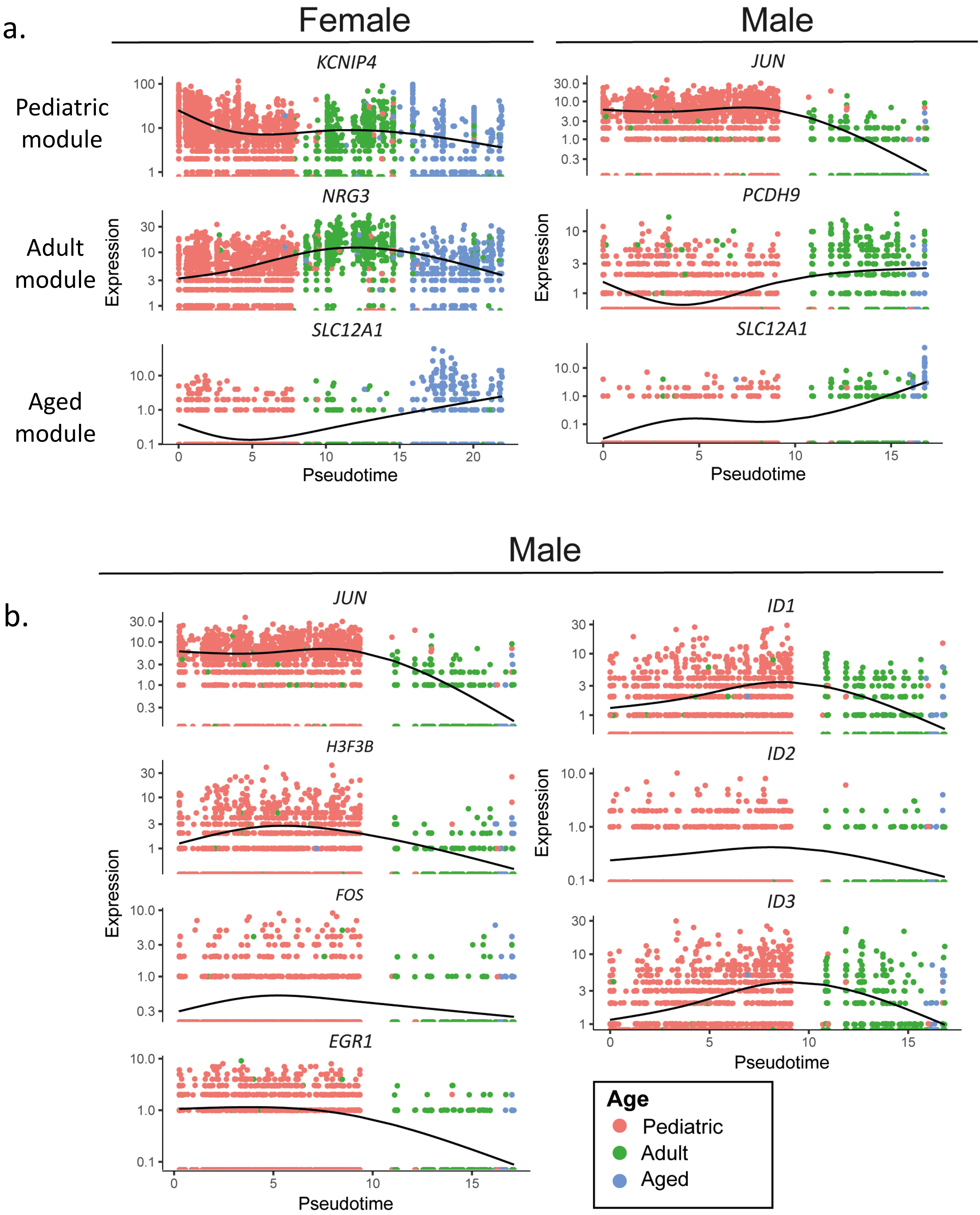


**Supplementary Figure 13:**

Analysis of the gene module dynamics for the human stem cell cluster. **a.** Trajectories of selected genes in females (*Left*) and males (*Right*) for the three age groups. **b**. Trajectories selected genes in the pediatric male pituitary sample, module 1. The color code for the ages of the subjects is presented.

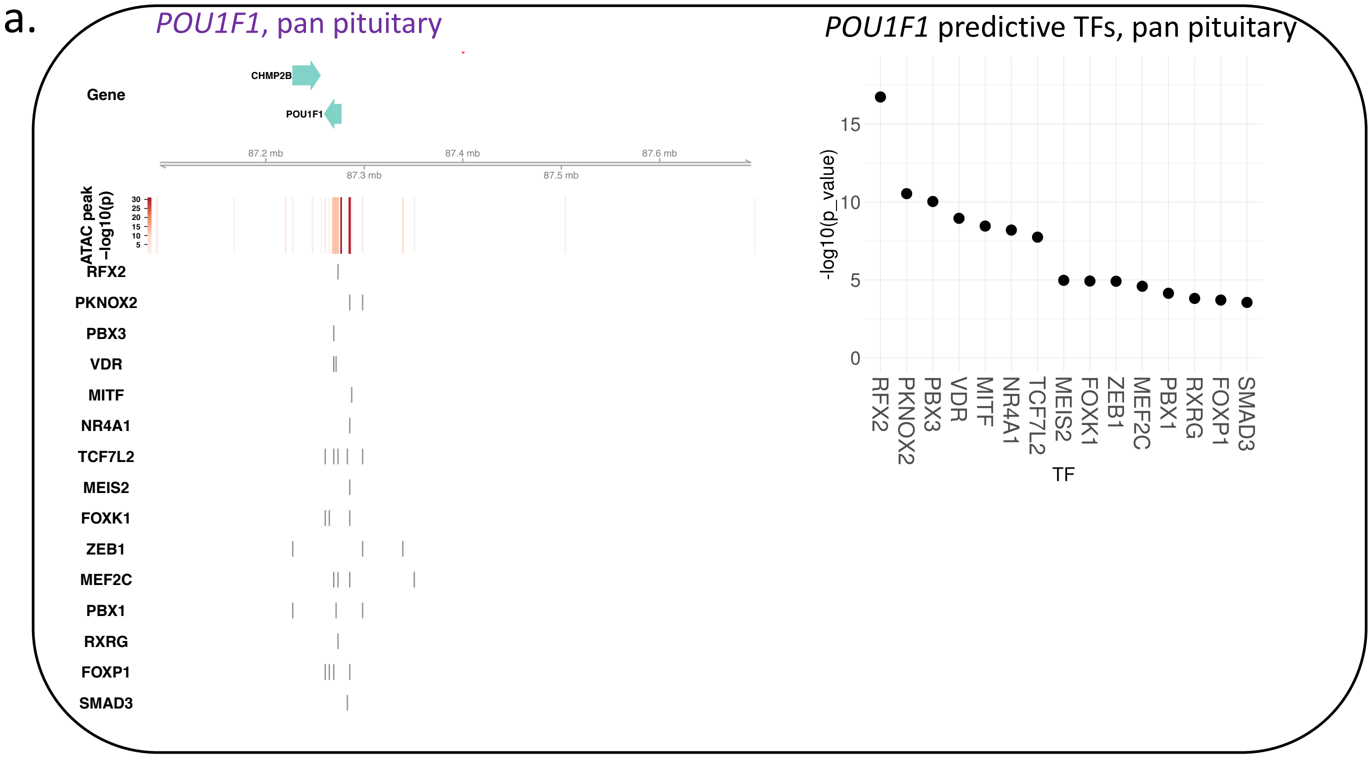


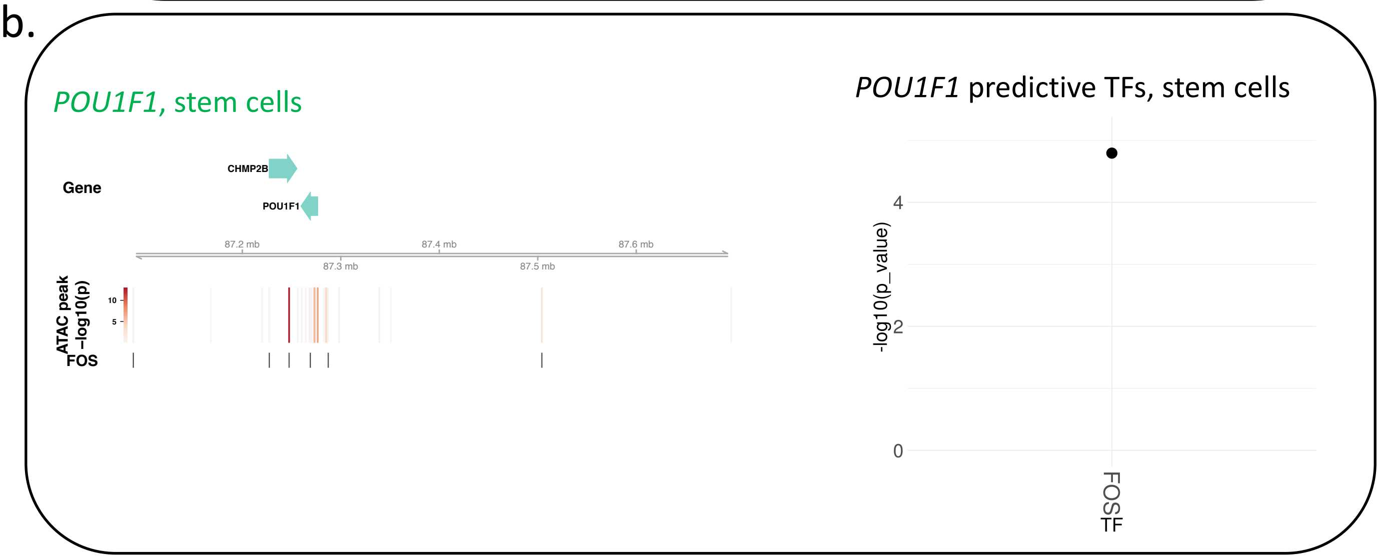


**Supplementary Figure 14: Linear model analysis identifies the main contributors for *POU1F1* specific gene expression in all pituitary cells vs. stem cells.**

**a**. Chromatin accessibility and the predicted TFs as modeled in all pituitary cells are shown for *POU1F1* expression (*Left*, top track is the contribution of each peak to gene expression measured by -log(p value), and bottom tracks are the TF binding sites). The individual contribution to *POU1F1* expression for the predicted TFs in all pituitary cell types is shown by -log(p value) (*Right*). **b.** Chromatin accessibility and the predicted TF for *POU1F1* expression is shown here as modeled in PSCs only (*Left*, top track is the contribution of each peak to gene expression measured by -log(p value), and the bottom track is the TF binding site). The individual contribution to *POU1F1* expression for the predicted TF in PSCs is shown as by -log(p value) (*Bottom*).

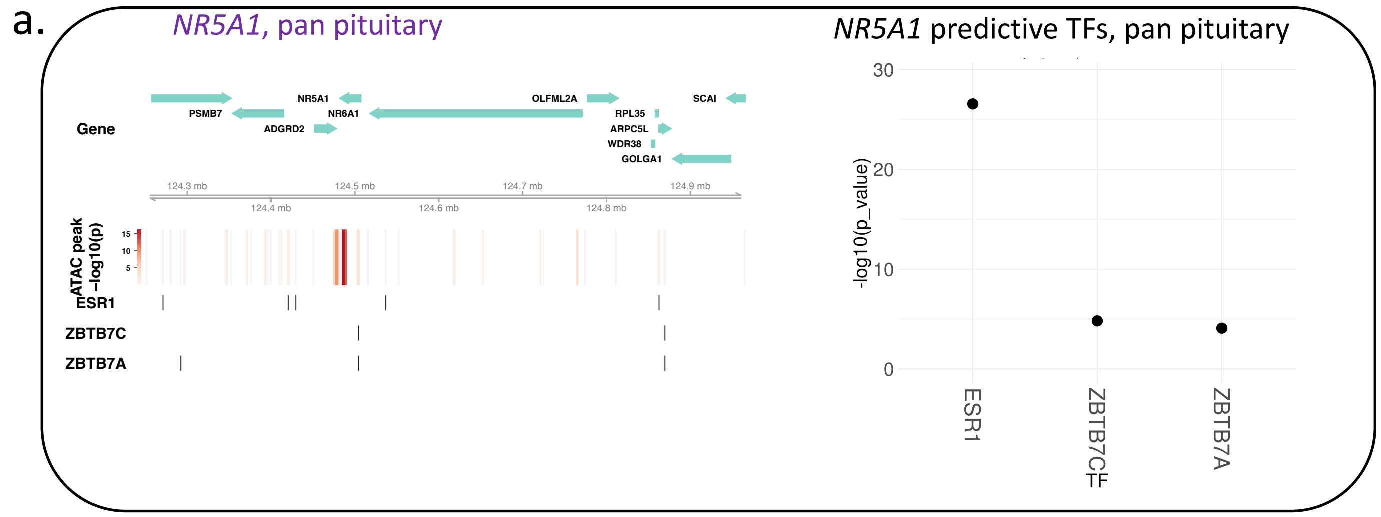


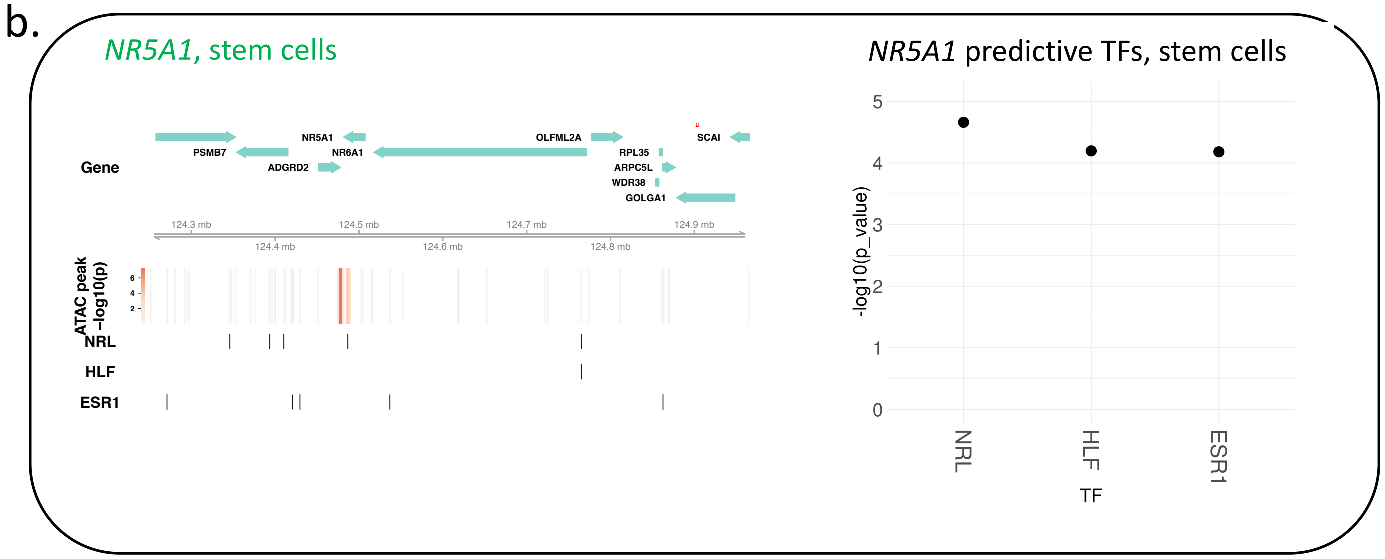


**Supplementary Figure 15: Linear model analysis identifies the main contributors for *NR5A1* specific gene expression in all pituitary cells vs. stem cells.**

**a.** Chromatin accessibility and the predicted TFs as modeled in all pituitary cells are shown for *NR5A1* expression (*Left*, top track is the contribution of each peak to gene expression measured by -log(p value), and bottom tracks are the TF binding sites). The individual contribution to *NR5A1* expression for the predicted TFs in all pituitary cell types is shown by -log(p value) (*Right*). **b.** Chromatin accessibility and the predicted TFs for *NR5A1* expression is shown here as modeled in PSC only (left panel, top track is the contribution of each peak to gene expression measured by -log(p value), and bottom tracks are the TFs binding sites). The individual contribution to *NR5A1* expression for the predicted TFs in PSCs is shown by -log(p value) (*Bottom*).

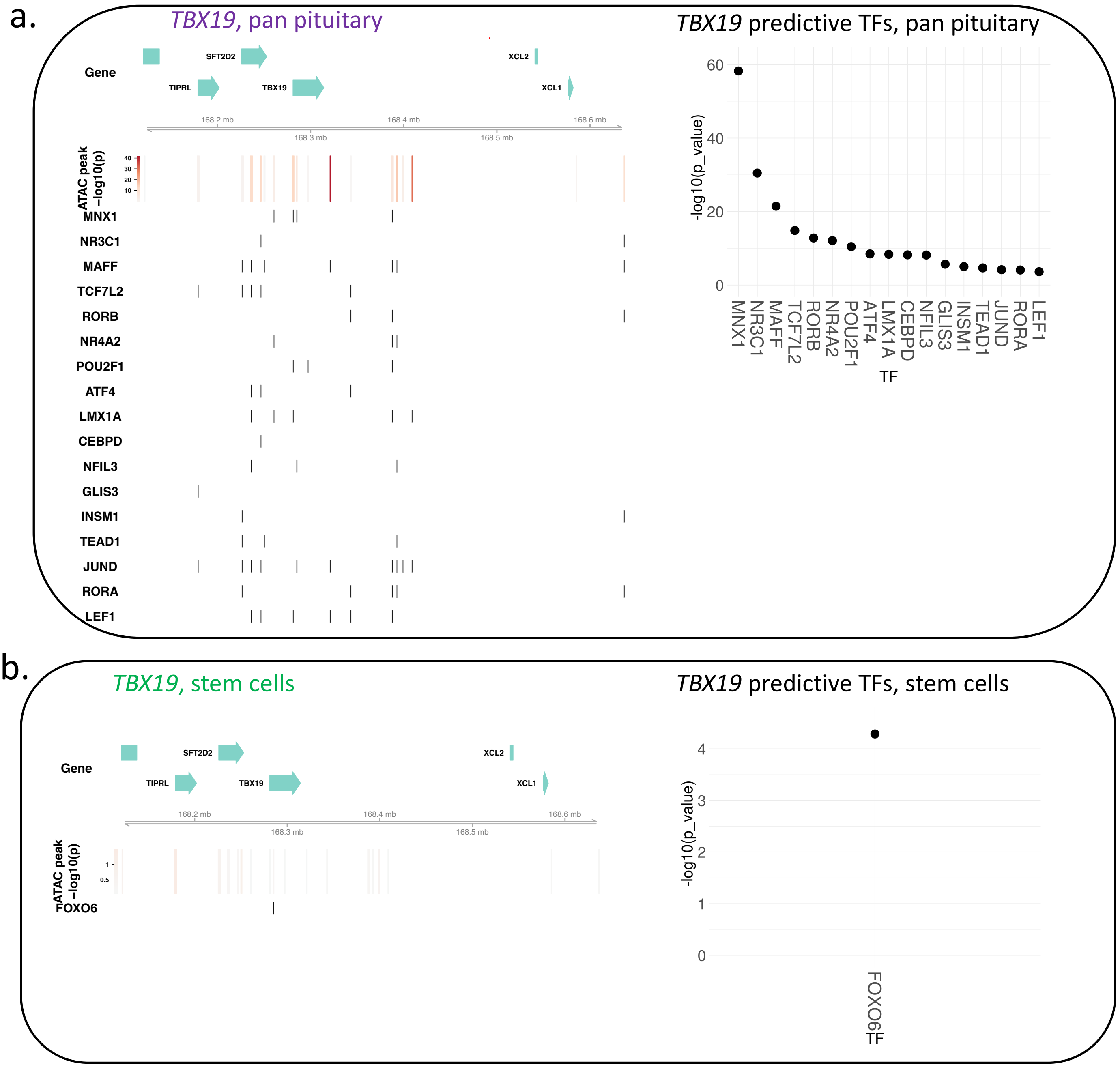


**Supplementary Figure 16: Linear model analysis identifies the main contributors for *TBX19* specific gene expression in all pituitary cells vs. stem cells.**

**a.** Chromatin accessibility and the predicted TFs as modeled in all pituitary cells are shown for TBX19 expression (*Left*,, top track is the contribution of each peak to gene expression measured by -log(p value), and bottom tracks are the TF binding sites). The individual contribution to *TBX19* expression for the predicted TFs in all pituitary cell types is shown by -log(p value) (*Right*). **b.** Chromatin accessibility and the predicted TFs for *TBX19* expression is shown here as modeled in PSC only (left panel, top track is the contribution of each peak to gene expression measured by -log(p value), and the bottom track is the TF binding sites). The individual contribution to *TBX19* expression for the predicted TF in PSCs is shown by -log(p value) (*Bottom*)

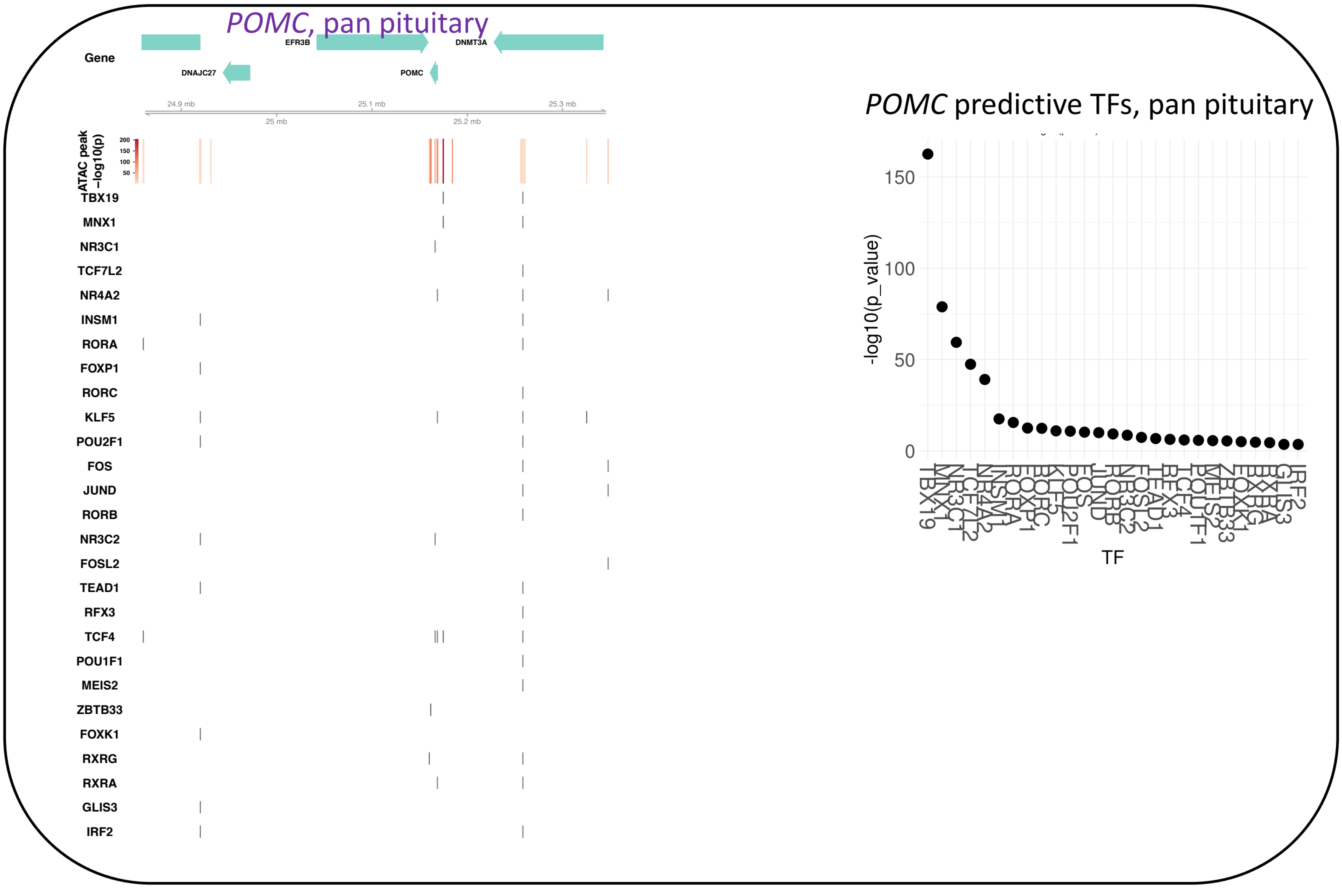


**Supplementary Figure 17: Linear model analysis identifies the main contributors for *POMC* specific gene expression in all pituitary cells**

Chromatin accessibility and the predicted TFs as modeled in all pituitary cells are shown for POMC expression (*Left*, top track is the contribution of each peak to gene expression measured by -log(p value), and bottom tracks are the TF binding sites). The individual contribution to *POMC* expression for the predicted TFs in all pituitary cell types is shown by -log(p value) (*Right*).

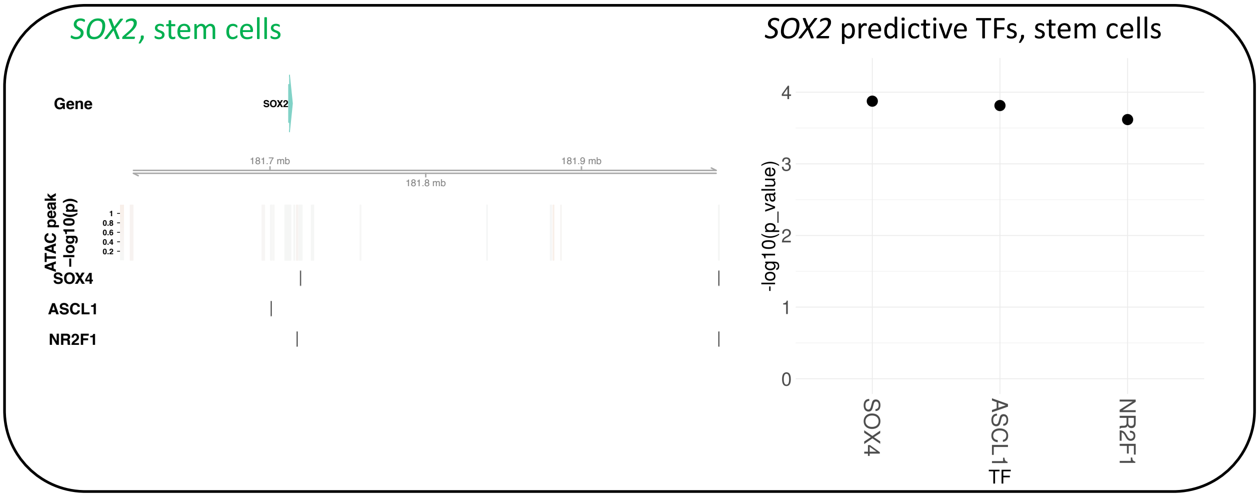


**Supplementary Figure 18: Linear model analysis identifies the main contributors for *SOX2* specific gene expression in stem cells**

Chromatin accessibility and the predicted TFs as modeled in stem cells are shown for SOX2 expression (*Left*, top track is the contribution of each peak to gene expression measured by -log(p value), and bottom tracks are the TF binding sites). The individual contribution to *SOX2* expression for the predicted TFs in stem cells is shown by -log(p value) (*Right*).
